# A reformulation of the *t*-test in single-cell transcriptomics dramatically reduces false positives

**DOI:** 10.1101/2025.03.12.642799

**Authors:** Oskar Kviman, Pedro F. Ferreira, Seong-Hwan Jun, Hosein Toosi, Jens Lagergren

**Author notes:** Contributing authors. These authors contributed equally to this work (Oskar Kviman, Pedro F. Ferreira).

## Abstract

The default differential expression tests in single-cell analysis software provide fast and simple approaches that help practitioners annotate cell populations identified through clustering algorithms. Here, we use theoretical arguments and empirical studies to show that these tests suffer from inflated false positive rates that affect the resulting annotations. We introduce LN’s ***t***-test, which tests a novel, asymptotically unbiased log-fold change estimator and avoids these false positives, without compromising speed.

## Main

Differential gene expression (DGE) analysis, the process of identifying the genes whose expression differs between groups of cells, is a fundamental ask in single-cell data analysis workflows [1]. This procedure is commonly used to annotate clusters of cells according to known marker genes or functional gene sets in an exploratory and iterative process [2, 3]. After clustering, DGE tests compare the expression levels in each cluster with the remaining population. Log-fold change (LFC) quantifies the relative difference in mean expression, and false positives are commonly defined as genes called differentially expressed but with negligible LFC [4] (Methods).

Standard single-cell analysis tools provide fast and simple tests to aid practitioners in cluster annotation: `scanpy` [5] defaults to a *t*-test, and `seurat` [6] defaults to the Wilcoxon rank-sum test.

While the LFC is the central quantity of interest in DGE testing, neither of these methods test it directly (Methods). Notably, the *t*-test, as implemented in both packages, applies the classical *t*-test framework to the difference in expectations of *log1p-transformed* random variables (Figure 1 a, b), and we refer to it as the log1p *t*-test. While the effect of the log1p transformation has been studied in the context of data normalization [7], and shown to create spurious differences between cells with different sequencing depths [8], its implications for DGE testing in cluster annotation have not been formally characterized.

**Fig. 1:**
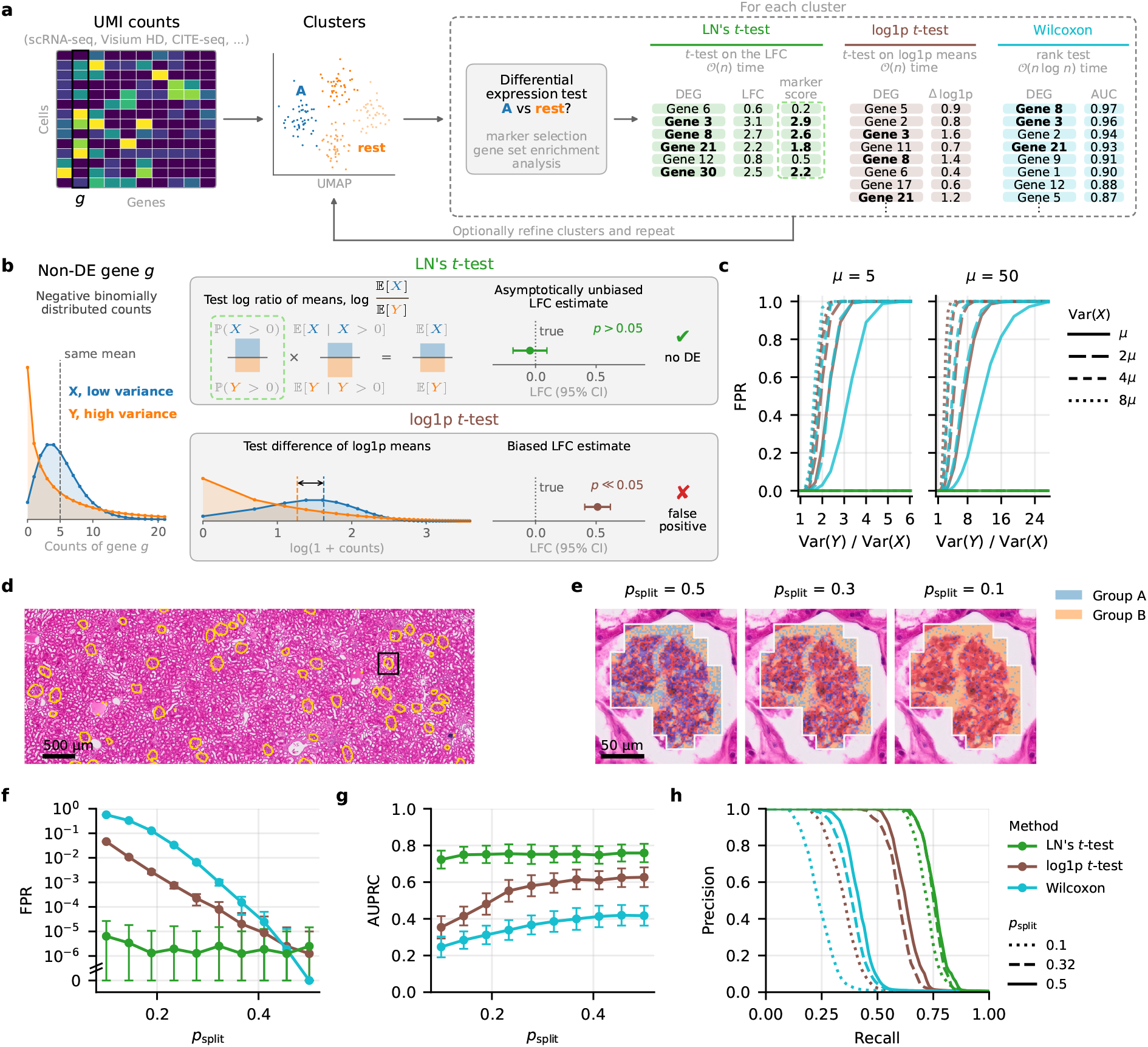
Variance heterogeneity induces biased differential expression that propagates to biological interpretation. **a**, The cluster annotation workflow. **b**, A gene with equal means but unequal variances in groups *X* and *Y* . LN’s *t*-test tests the log ratio of the means, whose estimate is asymptotically unbiased. The log1p transformation introduces mean differences that lead to false positives. **c**, FPR of each test on simulated groups *X* and *Y* with equal means *µ*. **d**, H&E image of a human kidney section (Visium HD), with the podocyte-containing glomerular capsules outlined. **e**, Spot splitting of a capsule into groups A and B, at three values of *p*_split_. **f**, FPR of each test without DEGs against *p*_split_. **g**, AUPRC of each test with simulated DEGs against *p*_split_. **h**, Mean precision–recall curves at *p*_split_ = 0.1, 0.32 and 0.5. Data in **f** –**h** are means over replicates, and error bars in **f** and **g** show standard deviations.

Here, we provide a theorem (Supplementary Note A.1) stating that, under negative binomial count data assumptions, the log1p *t*-test suffers from biased test statistics that result in false positives with probability 1 convergence whenever variance differs between groups (Figure 1 b, Methods), and provide a similar argument for MAST [9] (Methods). Because there is naturally a difference in gene expression variance between clusters – even for partitions of samples from a homogeneous population, as we show below – our theorem applies in practice. Additionally, as the Wilcoxon rank-sum test jointly tests differences in location and scale (Methods), variance imbalance renders its source of null rejection confounded.

To address these limitations, we developed LN’s *t*-test, a DGE method that factorizes the mean of a gene in each group into its detection rate and the mean of its non-zero counts (Figure 1 b). We approximate the estimator of each term with a log-normal (LN) distribution, and combine them into an asymptotically unbiased LFC estimator for DGE testing (Methods). For marker gene selection, LN ranks its DEGs by the log ratio of detection rates over their standard errors (Methods).

We first simulated negative binomial data with equal means and increasingly different variances (Methods), and found that LN’s *t*-test remained well calibrated, whereas the log1p *t*-test, Wilcoxon and MAST rapidly degenerated (Figure 1 c, Extended Data Figs 1 and 2), approaching a falsepositive rate (FPR) of one, as the theory predicts (Methods). LN’s *t*-test has the same complexity as the log1p *t*-test (Methods), and its runtime scaled linearly in the number of cells (Extended Data Fig. 2 c). The slow runtime of MAST is not compatible with the cluster annotation task, especially for large data sets.

Extending this controlled comparison to real data, we used Visium HD data of podocyte-containing glomerular capsules from healthy kidney tissue [10] (Figure 1 d, Supplementary Results 1), where each 2 *µ*m spot contains UMI counts. We randomly partitioned the spots within each capsule into one of two groups with probability *p*_split_ (Figure 1 e), and summed the counts of the spots assigned to each group. Decreasing *p*_split_ made the groups differ in sequencing depth and therefore increased variance differences, which led to an increased FPR for the log1p *t*-test, Wilcoxon and MAST, whereas LN’s *t*-test stayed below 10^−5^ (Figure 1 f, Extended Data Fig. 3 a,b). With synthetic differentially expressed genes (DEGs), LN’s *t*-test had the highest area under the precisionrecall curve (AUPRC) at every *p*_split_, with MAST approaching its performance only when the variances are balanced (Figure 1 g,h, Extended Data Fig. 3 c,d).

We then compared T cells with germinal center B cells from Visium HD data of a lymph node [11], taking the results from a pseudobulk DGE method as ground truth (Methods). We increased the variance imbalance without changing the biology by keeping each 2 *µ*m spot of the T cells with probability *p*_sub_ (Figure 2 a). LN’s *t*-test consistently recovered the ground truth on the T cells at a high AUPRC, while other methods degraded with decreasing *p*_sub_ (Figure 2 b; Supplementary Results 2). In gene set enrichment analysis (GSEA) of the MSigDB Hallmark gene sets [12] (Methods), the normalized enrichment scores (NES) obtained from LN’s gene ranking correlated highly with the pseudobulk result at every *p*_sub_, in contrast to the NES obtained from log1p *t*-test and Wilcoxon (Figure 2 c).

**Fig. 2:**
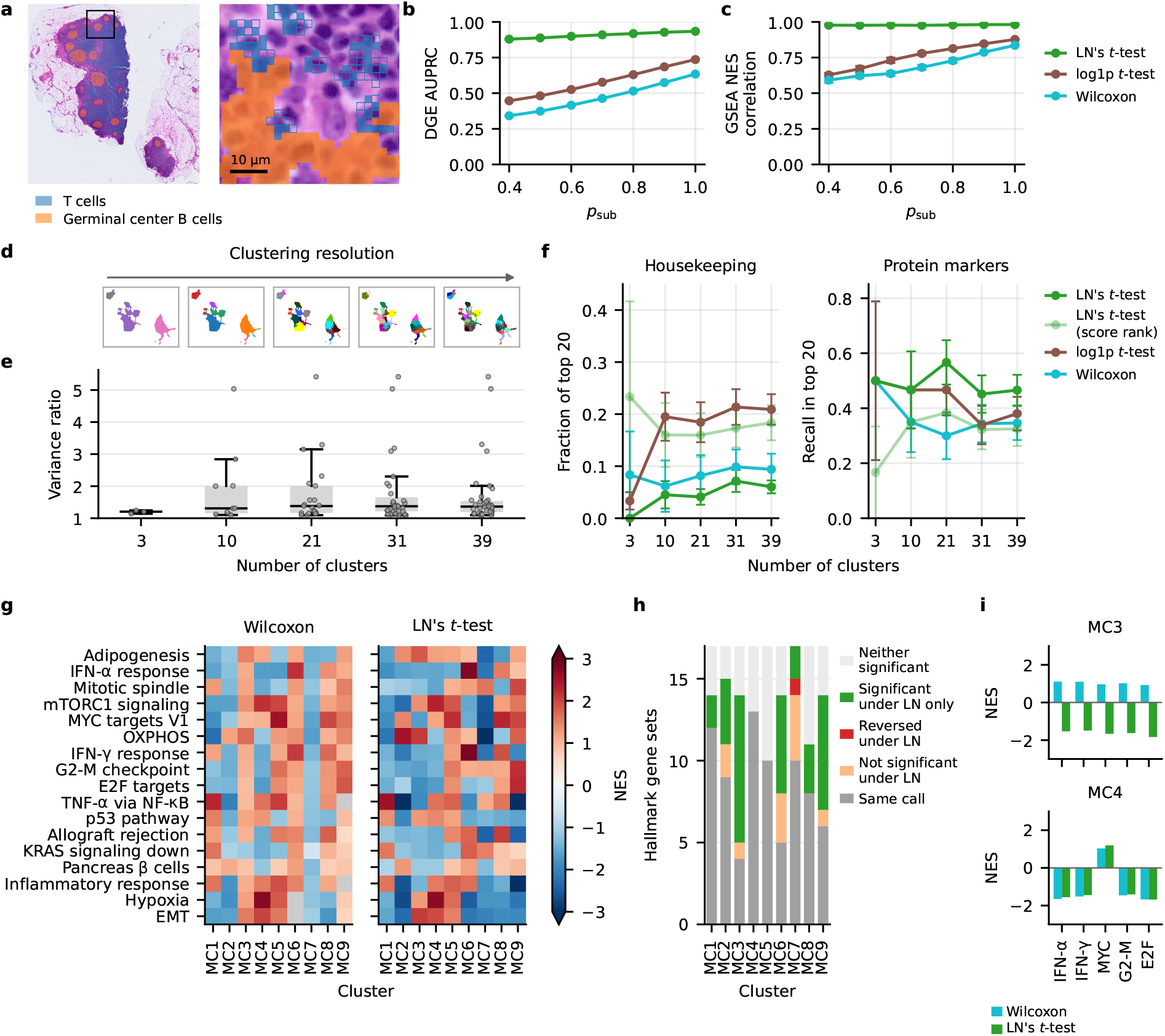
Depth-driven variance imbalances affect marker detection and pathway enrichments. **a–c**, Visium HD data from a human lymph node. **a**, H&E section and the boxed window, with the 2 *µ*m spots of T cells (blue) and germinal center B cells (orange). **b**, AUPRC of each test for the T-cell markers of the pseudobulk reference, against *p*_sub_. **c**, Agreement of each test’s Hallmark NES with the pseudobulk reference, against *p*_sub_. **d–f**, CITE-seq data from 10k PBMCs. **d**, UMAP coloured by Leiden clusters at five resolutions. **e**, Variance ratio between each cluster and the rest, against the number of clusters. **f**, Fraction of each test’s top 20 genes per cluster that are immune housekeeping genes [16] (left), and recall of the cluster’s surface-protein markers in the top 20 (right). For LN’s *t*-test (score rank), genes are ranked by test score instead of the marker score. **g–i**, scRNA-seq data from the glioma myeloid cell atlas [17]. **g**, NES of Hallmark gene sets for each cluster (MC1–MC9), with genes ranked by the Wilcoxon *z*-score (left) or LN’s score (right). Gray cells have no NES (Methods). **h**, For each cluster, the number of gene sets that keep their call under LN (same sign), lose it, reverse it, gain one, or are not significant under either test. Gray cells in **g** count as gained. **i**, NES of MC3 and MC4 in the interferon and proliferation gene sets. Data in **b** and **c** are means and standard deviations over replicates, and data in **f** are means and standard errors over the clusters of each resolution.

In CITE-seq and scRNA-seq data from PBMCs (PBMC10k [13] and PBMC3k [14]) clustered with Leiden [15] at five resolutions each, we observed variance imbalances between each cluster and the remaining cells (Methods, Figure 2 d,e, Supplementary Results 3). In PBMC10k, at every resolution, LN’s top 20 DEGs contained the fewest housekeeping genes [16] and recovered more of the cluster’s surface protein markers, which were measured in the same cells (Figure 2 f). The log1p *t*-test’s top 20 included the most housekeeping genes, and Wilcoxon recovered fewer protein markers. Ranked by test score instead of LN’s default marker score (Methods), the top 20 included more housekeeping genes and fewer protein markers, highlighting the benefit of LN’s LFC decomposition for marker gene selection. We also found that the LFCs that scanpy reports for the log1p *t*-test increased with resolution to a median of 15 in PBMC10k and 23 in PBMC3k, although the log1p means it tests differed by less than 0.6. Such discrepancies are expected because scanpy’s LFC approximation can greatly amplify small differences between transformed means (Supplementary Note I). Meanwhile, LN’s median stayed near 2 (Extended Data Fig. 4 c,d).

**Supplementary Figure 7:**
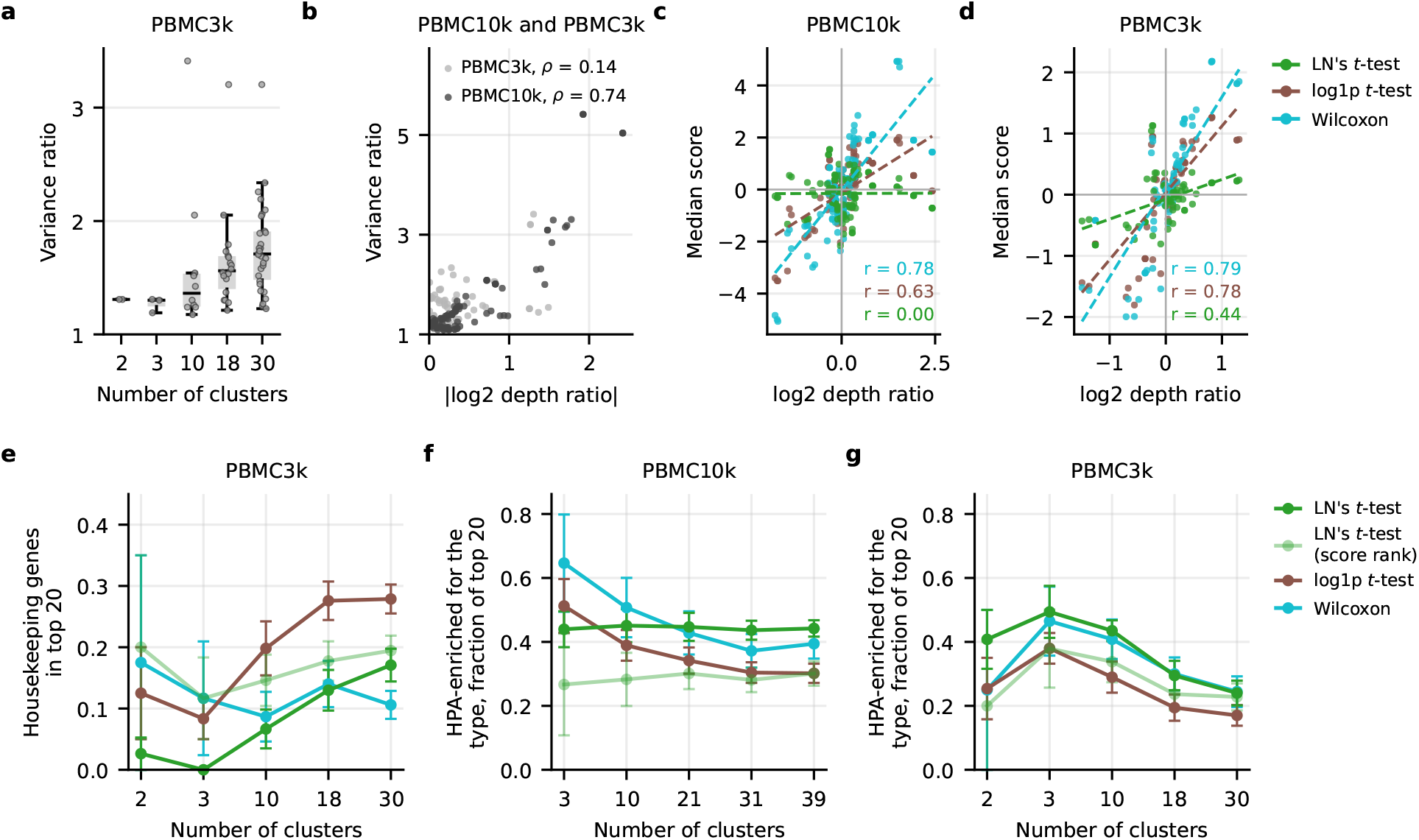
PBMC3k and PBMC10k multi-resolution clustering. **a**, Variance ratio between each cluster and the rest in PBMC3k. **b**, Variance ratio of every cluster against its |log_2_| depth ratio, in both data sets, with Spearman correlations. **c**,**d**, Median test score over each cluster’s expressed genes vs log_2_ depth ratios for (**c**) PBMC10k and (**d**) PBMC3k, with least-squares lines (dashed) and Pearson correlations. **e**, Fraction of the top 20 marker genes that are housekeeping genes, in the PBMC3k data. **f**,**g**, Fraction of the top 20 marker genes that overlap with the Human Protein Atlas [10] list for each cluster’s majority cell type, for (**f**) PBMC10k and (**g**) PBMC3k. Data in **e**–**g** are means and standard errors over the clusters of each resolution.

Finally, we applied LN’s *t*-test to a human glioma single-cell atlas [17]. We reproduced the workflow in the original study, which used GSEA on the DGE results from the Wilcoxon rank-sum test on nine myeloid cell clusters (Supplementary Results 4). Each cluster differed from the remaining myeloid cells in sequencing depth, which correlated with a variance imbalance that drove the Wilcoxon-based results (Extended Data Fig. 5). In contrast, LN was robust to this effect, resulting in different functional interpretations of some clusters (Figure 2 g,h). For example, the original study reported five pathways with contrasting enrichments in clusters MC3 and MC4, but LN found that the two clusters mostly agreed (Figure 2 i).

Our results demonstrate that LN’s *t*-test is robust to spurious variance imbalances in the groups under comparison, unlike the log1p *t*-test, Wilcoxon rank-sum test and MAST. LN’s *t*-test tests the LFC directly, providing directly interpretable effect sizes, at the computational cost of the log1p *t*-test. This makes it better suited for any application that requires fast and robust DGE testing, like cluster annotation and exploratory data analysis. LN’s *t*-test should therefore be the default for DGE analysis in `scanpy` and `seurat`.

Future extensions may address previously identified challenges in DGE testing, such as accounting for double-dipping (Methods) and batch effects [18], while retaining the computational speed that enables flexible exploratory data analysis.

## Methods

### 1 Notation

Fix a gene *g* and let *Y*_*g*_ and *X*_*g*_ denote the geneexpression random variables in the two groups (e.g. “cluster vs. rest” or “cluster *a* vs. cluster *b*”). We observe i.i.d. samples 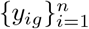 from *Y*_*g*_ and 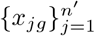 from *X*_*g*_, where *y*_*ig*_, *x*_*jg*_ ∈ ℝ_≥0_ are (optionally depth-normalized) counts. We denote *G* as the total number of genes, and log1p(*x*) = log(*x* + 1).

### 2 Why test LFC?

While we are considering an LFC-based null hypothesis in this work, expression variability can carry phenotypic information for biological discovery [19], and differential variability may indeed complement LFC-based analysis [20].

However, it is important to distinguish changes in variability from changes in mean expression. A significant result that can arise from either does not, by itself, establish that a gene is up- or downregulated. Notably, Wilcoxon rejects the null based on differences in location and/or shape [21], and it has long been disadvised to use it as a test for differences in locations [22] (more on Wilcoxon below and in Section H).

Resolving the ambiguity of which expression feature triggered the rejection requires further analysis, adding a step each time differential expression is assessed. For iterative clustering and annotation pipelines that require efficient identification of differentiated genes, an LFC-based null therefore provides a direct match between the statistical hypothesis and the biological interpretation: significance supports a change in mean expression, while the LFC quantifies its direction and magnitude.

As we have shown (Theorem 1), the log1p *t*-test is a biased test of mean differences that accidentally, under an equal mean condition, instead rejects the null due to variance imbalance. Unfortunately, the effect size does not have an easy analytical form due to the log1p transform, and so any significance measured by the log1p *t*-test is, as for Wilcoxon, a result of confounding factors: was there a difference in means/location or variance/scale? Meanwhile, LN’s *t*-test unequivocally tests difference in LFC via a new asymptotically unbiased LFC estimator (details below).

### 3 Time complexity analysis

The time complexity of the log1p and LN’s *t*-test is *O*(*G*(*n* + *n*^*′*^)). Meanwhile, due to its ranking step, the time complexity of the Wilcoxon rank-sum test is *O*(*G*(*n* + *n*^*′*^) log(*n* + *n*^*′*^)).

### 4 The log1p-transform biases DGE tests

Under the typical assumption that these random variables follow negative binomial (NB) distributions with equal mean, i.e. E[*Y*_*g*_] = E[*X*_*g*_] but different dispersions, we prove (Lemma 7) that the difference in mean of the log1p-transformed (optionally depth-normalized; Remark 2) random variables is biased in the sense that

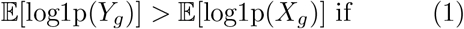

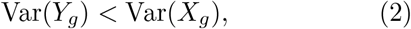

and vice versa if Var(*Y*_*g*_) > Var(*X*_*g*_). See Supplementary Note G for an intuition of how the difference in variance directly affects the difference in mean of the log1p-transformed random variables.

#### 4.1 *t*-tests on log1p-transformed samples inflate FPRs

In the *t*-test implementations in both `scanpy` and `seurat`, the effect size is the difference in the empirical mean estimates of the log1p-transformed samples, i.e. the difference between the following two estimators

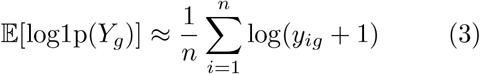

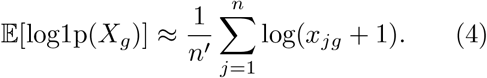

These unbiased estimators converge to their target expectations, which results in biased *t*-tests for sufficiently large sample sizes (Theorem 1). Moreover, the log1p *t*-test is also biased for finite sample sizes but sufficiently large variance differences Conjecture 1).

### 4.2 MAST will produce high FPRs

Here we demonstrate that our theoretical results also apply to MAST in an unadjusted two-group comparison [23], and thus that it produces large FPR scores under the LFC-based null hypothesis and variance imbalance.

Let 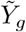 denote the log1p-transformed, CP10K-normalized version of the gene expression r.v. *Y*_*g*_ i.e., the input to MAST). To model gene expresion distributions, MAST uses a *hurdle model* with logistic regression to model the detection probability through the logit link

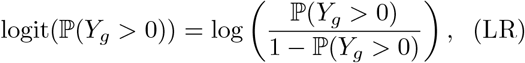

and, following a normality assumption on 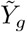, Gaussian linear regression on the mean of 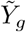 for *g* with non-zero UMI-counts, i.e.

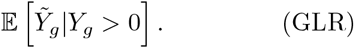

MAST then constructs two non-negative test statistics that evaluate group-difference in LR and GLR, respectively, before adding the statistics and comparing their sum to the null hypothesis.

Dropping the gene subindex, let *p*_*Z*_ = ℙ(*Z* > 0) and *a*_*Z*_ = E[log(1 + *Z*) *Z* > 0], for *Z X, Y* . For negative binomial variables with equal means and Var(*X*) < Var(*Y*), Lemma 7 gives

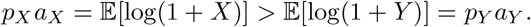

Consequently, *p*_*X*_ = *p*_*Y*_ and *a*_*X*_ = *a*_*Y*_ cannot both hold: the null hypothesis is false w.r.t. at least one of MAST’s two test statistics. Since the test ums the non-negative component statistics, evidence against either null cannot be canceled by the other component. In a two-group model with independent observations, the joint test therefore ejects with probability approaching one as both ample sizes grow. These rejections occur despite equal mean expression and thus MAST systemat-cally calls false positives under our definition of differential expression. In Extended Data Fig. 2 we show how MAST suffers identically to the log1p *t*-test and Wilcoxon on synthetic data due to variance imbalance, and occasionally worse than all other methods on real data in Extended Data Fig. 3. Meanwhile, LN’s *t*-test is robust, as it only tests for LFC differences.

### 5 Wilcoxon’s high FPRs are expected

We provide a formal explanation of why the Wilcoxon rank-sum test calls a gene DE when the two populations have the same mean but different variances, in Supplementary Note H. Here, we emphasize that this behavior is expected based on the statistics literature: the Wilcoxon ranksum test, or, equivalently, the Mann-Whitney *U* -test, is a test of difference in distributional location and/or shape [21]. In fact, the test only measures a difference in location (e.g. median) under a highly restrictive assumption on equal distribution shapes (e.g., variance; [22]), although scientific misuse of the test has been recorded over the course of two decades [22, 24]. Instead, the Wilcoxon rank-sum test tests whether samples from one group tend to be larger than those from the other group, rather than focusing on differences in averages. When the data are non-negative, as in expression data, the Wilcoxon rank-sum test will thus call genes with equal mean but difference in population variance differentially expressed. As we have shown, in cluster annotation tasks, difference in variance between clusters naturally occur without any meaningful biological signal, e.g. between partitions of samples from an homogeneous population. Thus, the striking numbers of observed false positives that the Wilcoxon rank-sum test output are expected.

### 6 LN’s *t*-test

LN’s *t*-test is a differential expression test that applies the classical *t*-test framework to a novel effect size estimator that is an asymptotically unbiased estimator of the log-fold change (LFC).

Unlike the log1p *t*-test, LN’s *t*-test does *not* test for differences in expectations of log1p-transformed random variables. Instead, it targets the LFC between the population means, i.e. the target estimand is the (base-2) log-fold change

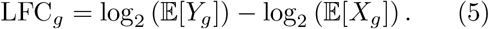

See Supplementary Note B for an in-depth derivation of the method.

#### 6.1 Mean factorization

LN’s *t*-test is based on the identity

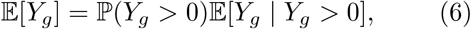

and analogously for *X*_*g*_. Hence, inference on LFC_*g*_ reduces to estimating detection probabilities and conditional means in each group. The two subsequent sections are devoted to developing LN distributed estimators of these two terms. By doing so, we obtain an LFC estimator that is normally distributed with unbiasedness guarantees, and closed-form confidence intervals.

#### 6.2 Estimating detection probability

Define the number of detected (non-zero) samples

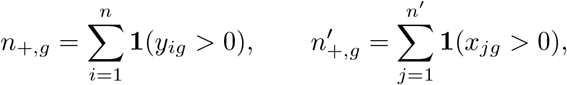

and the empirical detection rates

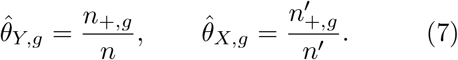

To propagate uncertainty on the log scale, we approximate 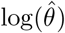 using a Beta moment model:

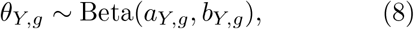

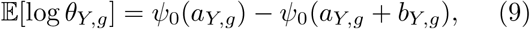

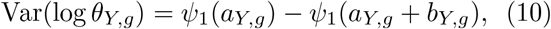

where *ψ*_0_ and *ψ*_1_ are matched, truncated largeargument asymptotic expansions of the digamma and trigamma functions, r espectively [25, Eq. 5.11.2 and 5.15.8]:

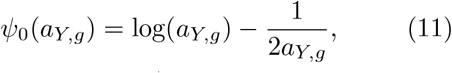

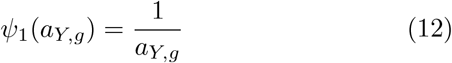

(and analogously for *θ*_*X,g*_). Their first-order correction terms cancel in the induced log-normal mean, ensuring that it exactly matches the Beta mean *a/*(*a* + *b*) (Equation (B15)).

Using these expressions for the mean and variance of log(*θ*), and replacing *a*_*Y,g*_ *b*_*Y,g*_ with *â*_*Y,g*_ = *n*_+,*g*_ and 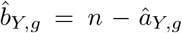 (vice versa for *a*_*X,g*_), we parameterize a log-normal (LN) approximation of the Beta pdf on the natural scale, such that 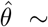 LN (E[log *θ*], Var(log *θ*)). (See Supplementary Note B.1 for details and an unbiasedness proof.) Interestingly, we believe that this is the first recorded LN approximation of a Beta pdf.

#### 6.3 Estimating the conditional mean

Let *π*_*Y,g*_ = {*i* : *y*_*ig*_ > 0} and *π*_*X,g*_ = {*j* : *x*_*jg*_ > 0}. Define the sample mean among detected cells

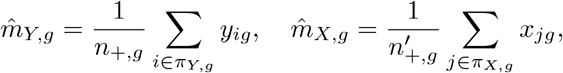

and 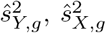 as the corresponding sample variances. We model 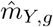 as log-normal with estimated parameters

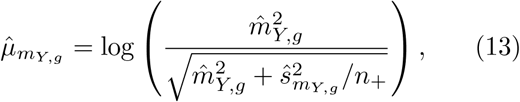

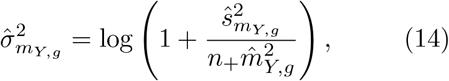

and note that the estimator is clearly unbiased: 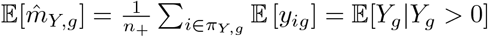.

#### 6.4 LN LFC estimator and standard error

We define the LN effect size estimator

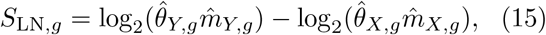

and prove that it is an unbiased estimator of LFC_*g*_ under mild conditions (Theorem 2). Under the log-normal approximations, *S*_LN,*g*_ is approximately normal (Lemma 2) with variance given by the sum of log-variances. Concretely, writing log_2_(*z*) = log(*z*)*/* log(2), we use the standard error

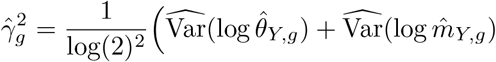

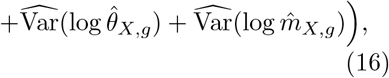

where 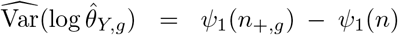 and 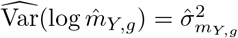.

#### 6.5 Test statistic, *p*-values, and confidence intervals

We test

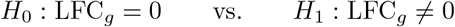

using the *t*-test-style statistic

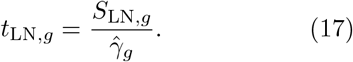

We report two-sided *p*-values based on the standard *t*-testing framework, and statistics of the *t* distribution were computed using `scipy.stats.distributions.t`. Uncorrected significance is calculated using the (1 − *α*) confidence interval

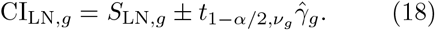

#### 6.6 Multiple testing correction

For gene-wise testing over *G* genes, we correct *p*-values using Bonferroni unless otherwise stated. We used Bonferroni in an attempt to salvage he FPR scores of the baseline methods, as it is more conservative. Genes are called differentially expressed at FPR level *q* if the Bonferroniadjusted *p*-value is below *q*.

#### 6.7 Marker score

A cluster’s markers are the genes LN’s *t*-test calls significant with a positive statistic, ranked by heir marker score: with the cluster as *Y* and the remaining cells as *X*, the log ratio of the detection rates (the detection term of (6)) over its standard error,

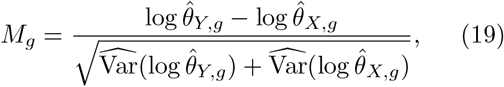

with the variances of (16) and *n*_+,*g*_ and 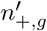 floored at 1; ties are broken by *t*_LN,*g*_, then by gene name.

### 7 Regarding double-dipping

Double-dipping in single-cell transcriptomics [26] affects all DGE methods that hypothesis test with a cluster-dependent null hypothesis. I.e., a null hypothesis that states that there is no cluster-level difference between gene expression test targets (e.g. distribution means or variability), although the clusters are partitions of the data based on their differences in gene expression patterns. As an informal, illustrative example of this mechanism, consider the following setup.

We observe and partition samples from a single, homogeneous population into two clusters. Then, the clusters are by design grouped into partitions merely due to variance in the sampling. Trivially and meaninglessly, there will be a detectable difference in means and/or variance between these two clusters, and so we can expect a large fraction of false positives.

Double-dipping is an important consideration for inference after clustering. Count splitting provides a framework for separating clustering from downstream testing, with developments for both Poisson [27] and negative binomial data [28]. Incorporating such approaches into workflows using LN’s *t*-test is an important direction for future work, complementary to our analysis of how the choice of test target affects DGE testing.

### 8 DGE method settings

We ran LN’s *t*-test with the `ln-ttest` Python package on counts normalized to 10,000 per cell (CP10K). The log1p *t*-test and the Wilcoxon rank-sum test were run with `scanpy’s rank genes groups` [29] on log1p-transformed CP10K, using tie correction for Wilcoxon. For MAST [23], we used `FindMarkers` from the seurat R package with its gene filters turned off (`logfc.threshold = 0, min.pct = 0`), so that every gene is tested, as for the other methods. DEGs were called at Bonferroni-adjusted *p* < 0.05. The effect sizes in Extended Data Fig. 4 are the ones reported by each package, i.e. LN’s LFC estimate, `scanpy`‘s `logfoldchanges` for the log1p *t*-test and `seurat`‘s `avg log2FC` for Wilcoxon. Note that `scanpy`‘s `logfoldchanges` is the log_2_ of the ratio of exp(·) − 1 of each group’s mean log1p CP10K (Supplementary Note I), which is not the effect size that the log1p *t*-test tests. We therefore also show the difference in mean log1p CP10K.

### 9 Evaluation metrics

FPR is the fraction of non-DEGs that a method calls as DEGs, and TPR is the fraction of true DEGs called as such. AUPRC is the average precision of the genes ranked by each test. To compare variances at equal mean, we computed the variance ratio between two groups on CP10K over the genes detected in at least 5 cells of each group and whose |log_2_| mean ratio is at most 0.1, i.e. genes with approximately equal means, as 2 to the power of the median |log_2_| variance ratio. The depth ratio of a group is its median counts per cell over that of the other group. A gene is considered expressed if it is detected in at least 5% of either group’s cells. Error bars show the mean *±* s.d. over replicates, or *±* s.e.m. over clusters.

### 10 Synthetic data

Counts were drawn from negative binomial distributions with mean *µ* and variance *µ* + *ϕµ*^2^ (Poisson for *ϕ* = 0). For the experiments without DEGs, we simulated two groups of 1000 cells with a shared mean *µ* = 5 or 50 across 1500 genes, with Var(*X*) = *µ*, 2*µ*, 4*µ* or 8*µ* and Var(*Y*)*/*Var(*X*) varying from 1 to 64. In the simulations including MAST, only 150 of the 1500 genes of *Y* took the larger variance, while the rest were identical in both groups. This way, the detection rates of the cells barely differ between the groups, as when only a subset of genes changes in real data. Here, we used Var(*X*) = *µ* or 4*µ* and ratios from 1 to 32. The FPR was computed over the 150 genes. For the LFC intervals, we simulated two groups of 1000 cells and 1500 genes. We set *X* with *ϕ* = 2 and mean 10, with a tenth of its genes shifted up by |*ϵ*|, *ϵ* ∼ *N*(0, 5), and *Y* with *ϕ* = 1 and mean 10*e*, i.e. *e* times deeper, so that the true LFC after depth normalization is log_2_(10*/µ*_*X*_). For the simulations with DEGs, we simulated two groups of 10000 cells and 1500 genes in 20 replicates. Each gene was a DEG with probability 0.1, in which case its mean in *X* was multiplied or divided by 1 + *s/µ* with equal probability. In the sparse regime, we set the non-DEG mean to 10, *s* = |*ϵ*| with *ϵ* ∼ *N* (0, 5), *ϕ*_*Y*_ = 1 and *ϕ*_*X*_ ∈ {1, 1.5, 2}. In the dense regime, we set the mean to 100, *s* = |*ϵ*| with *ϵ* ∼ *N* (15, 5), *ϕ*_*Y*_ = 0.1 and *ϕ*_*X*_ ∈ {0.1, 0.2, 1}. Finally, we measured runtimes on one thread, on sparse counts of 2000 genes with log *µ* − *N*(log 0.15, 1.3^2^) and *ϕ* = 0.5, for 1000 to 100000 cells split equally into two groups. We report the median of three runs.

### 11 Kidney Visium HD data

We used the Visium HD human kidney data from [30]. Glomerular capsules were taken as the connected regions of the 16 *µ*m bins in Space Ranger’s cluster 5, and every 2 *µ*m spot inside a capsule was assigned to it. This resulted in 161 capsules with 18085 detected genes. As a negative control, we split the capsules into two groups at their median total UMI count, first with all 161 capsules and then removing the smallest capsules. All other kidney experiments removed the 35 capsules with the fewest UMIs, leaving 126 capsules. In the spot splitting experiment, we assigned the spots of each capsule to group A with probability *p*_split_ and to group B otherwise. We then summed the spots of each group into one profile per capsule. This creates two groups from the same capsules that differ in sequencing depth whenever *p*_split_≠ 0.5. We varied *p*_split_ in 10 equal steps from 0.1 to 0.5, with 100 replicates each. Owing to its runtime, MAST was only run on the first 3 replicates of each *p*_split_, and in Extended Data Fig. 3 a,b the other tests are shown on the same 3 replicates. In the UMI downsampling experiment, we thinned the counts of every capsule binomially with probability *p*_sample_ and then split the capsules at their median depth. Without DEGs, we varied *p*_sample_ in 10 equal steps from 0.1 to 1, with 10000 replicates each. With DEGs, we used 10 equal steps from 0.125 to 1, with 100 replicates each. In both experiments, we introduced DEGs by selecting each gene with probability 0.005 and multiplying its counts by 2^3^ in a randomly chosen group. Genes with no counts in one of the two groups were dropped before testing.

### 12 Lymph node Visium HD data

We used the Visium HD human lymph node data [31] with the cell segmentation from 10x Genomics, assigning every 2 *µ*m spot to its segmented cell and every cell to its Leiden cluster. We hen compared the T-cell zone (cluster 1) with the germinal center B cells (cluster 3), keeping cells hat had at least 3 spots and 20 UMI counts, and genes detected in at least 2 cells in each group. As ground truth, we built a pseudobulk reference on he full data. Here, we shuffled the cells of each group and summed them into pools of 200 cells, which served as the samples for a Wald test with PyDESeq2 [32], with Benjamini–Hochberg correcion over all genes. Genes with adjusted *p* < 0.05 were taken as the markers of the group in which hey have a positive LFC. To lower the sequencng depth of the T cells without changing their biology, we kept each of their 2 *µ*m spots with probability *p*_sub_ from 0.4 to 0.9 in steps of 0.1. In a second experiment, we instead kept each T-cell UMI with probability *p*_sample_ from 0.2 to 0.9 in teps of 0.1. Each *p*_sub_ and *p*_sample_ had 20 replicates. For GSEA, all tested genes were ranked by each test’s score, germinal center B cells against T cells, and by the PyDESeq2 Wald statistic. We hen ran GSEApy’s prerank [33] on the MSigDB Hallmark sets [34] with 1000 permutations on the first 10 replicates. Finally, we computed the Spearman correlation between the NES of each test and hose of PyDESeq2.

### 13 PBMC data

PBMC10k is the RNA of 5’ CITE-seq run on healthy-donor PBMCs from 10x Genomics [35]. We kept cells with at most 10% mitochondrial counts and at least 500 antibody counts, and removed doublets with Scrublet [36]. PBMC3k is he 3k PBMC data set from 10x Genomics [37]. In both data sets, we removed cells with fewer han 200 genes and genes detected in fewer than 3 cells. For clustering, we used the 2000 most variable genes (`Seurat` v3 flavour), log1p CP10K, scaling, 40 principal components and a 10-nearestneighbour graph, with Leiden [38] at resolutions 0.01, 0.1, 1, 2 and 3. Each cluster was then tested against all other cells on all genes. Cells were labelled with CellTypist’s [39] Immune All Low model with majority voting, and each cluster takes the majority type of its cells. Housekeeping genes are the immune housekeeping genes of Monaco et al. [40]. Cell-type-enriched genes are those that the Human Protein Atlas (release 25.1) [41] lists as immune cell enriched, or as group enriched with the cluster’s type in the group. In PBMC10k, a cluster’s protein markers are the genes of the 13 single-gene antibodies whose centred log-rationormalized level separates the cluster from the rest with an AUC of at least 0.75. We computed recall as the fraction of each cluster’s protein markers in the top 20.

### 14 Human glioma atlas data

We used the count matrices (GEO GSE182109) and cell metadata of the glioma myeloid cell atlas [42], which contain 83479 myeloid cells in nine clusters. We kept the genes detected in at least 3 cells. Each cluster was tested against the other myeloid cells on log1p CP10K. As in the code of the original study, we ranked genes by presto’s [43] uncentred Wilcoxon AUC and ran fgsea [44] on the 50 MSigDB Hallmark sets [34] (msigdbr 7.5.1) with minimum set size 10 and its default multilevel method, with 10^4^ permutations for the initial estimate (10^5^ when a set was left without an NES). We then compared the signs of the resulting NES with the enrichment directions reported in the original study. For the comparison between LN’s *t*-test and Wilcoxon, we ranked genes by their test statistics, i.e. LN’s score and scanpy’s tie-corrected Wilcoxon *z*-score. A gene set is called at fgsea’s adjusted *p* < 0.05. For four cluster-gene set pairs, the *z*-score ranking had no NES, because no random set took the observed sign even with 10^5^ permutations. DEGs were called at Bonferroni-adjusted *p* < 0.05 over 24257 genes. Finally, the UMAP was computed with seurat and Harmony [45], as described in the original study.

## Data availability

The synthetic data were generated by code available at https://github.com/okviman/lntest. The Visium HD human kidney data [30], the Visium HD lymph node data [31], the PBMC10k data [35] and the PBMC3k data [37] were retrieved from the publicly available 10x Genomics website, and the human glioma data are available from [42]. The Hallmark gene sets are from MSigDB [34], the immune housekeeping genes from Monaco et al. [40] and the cell-type marker genes from the Human Protein Atlas (release 25.1) [41].

## Code availability

LN’s *t*-test is available as the Python package `ln-ttest` at https://github.com/okviman/lntest, which also contains the code that reproduces every analysis and figure presented here.

## Extended Data

**Extended Data Figure 1:**
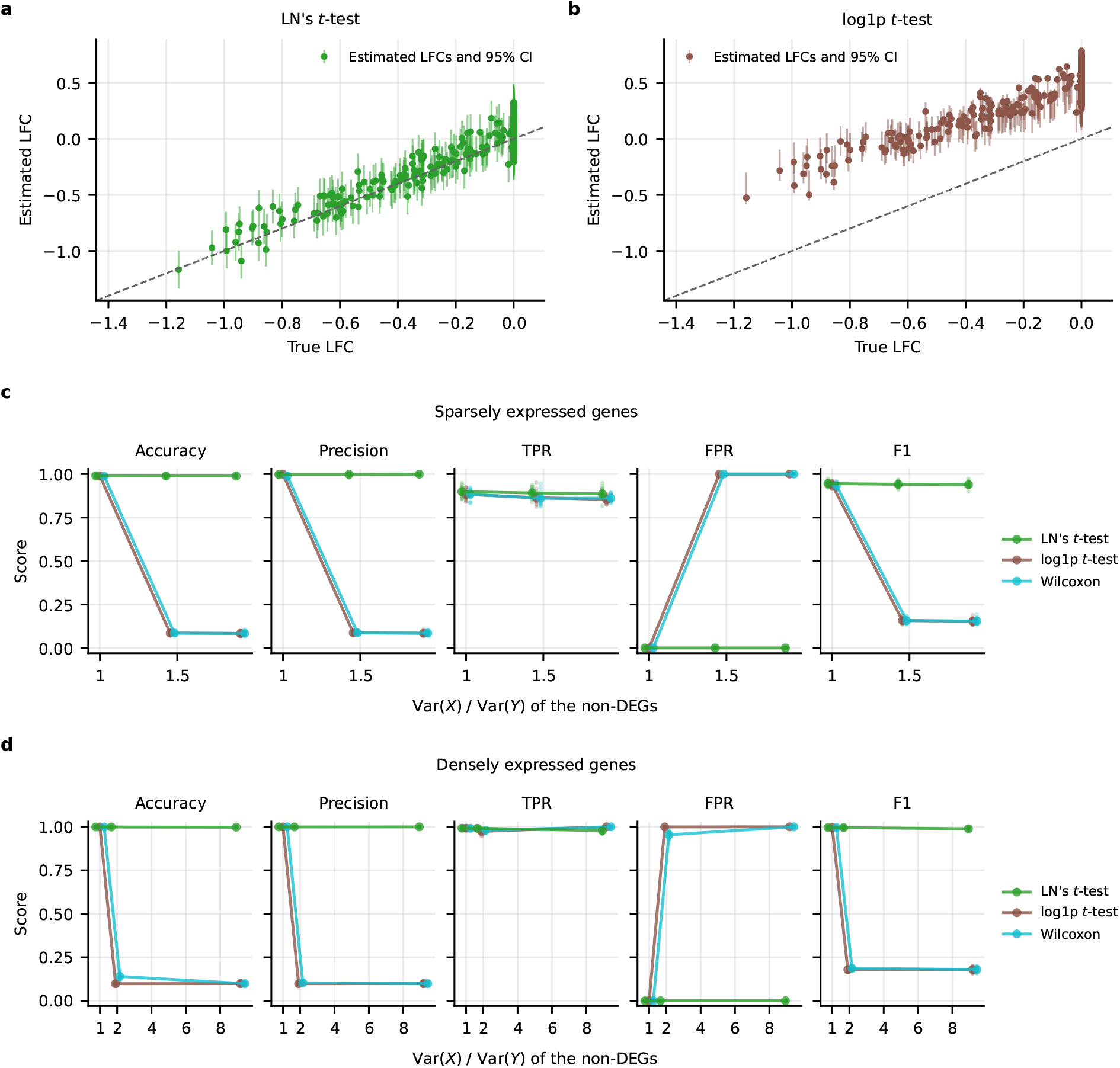
Synthetic negative binomial data. **a**,**b**, Estimated log-fold change (LFC) with its 95% confidence interval against the true LFC, for LN’s *t*-test (**a**) and the log1p *t*-test (**b**). For the log1p *t*-test, the points are the LFCs that `scanpy` reports and the intervals are those of its test. The dashed line marks equality. **c**,**d**, Accuracy, precision, true-positive rate (TPR), false-positive rate (FPR) and F1 of each test with simulated differentially expressed genes (DEGs), against the variance ratio Var(*X*)*/*Var(*Y*) of the non-DEGs, for sparsely (**c**) and densely (**d**) expressed genes. In **c** and **d**, dots are replicates and lines are their means.

**Extended Data Figure 2:**
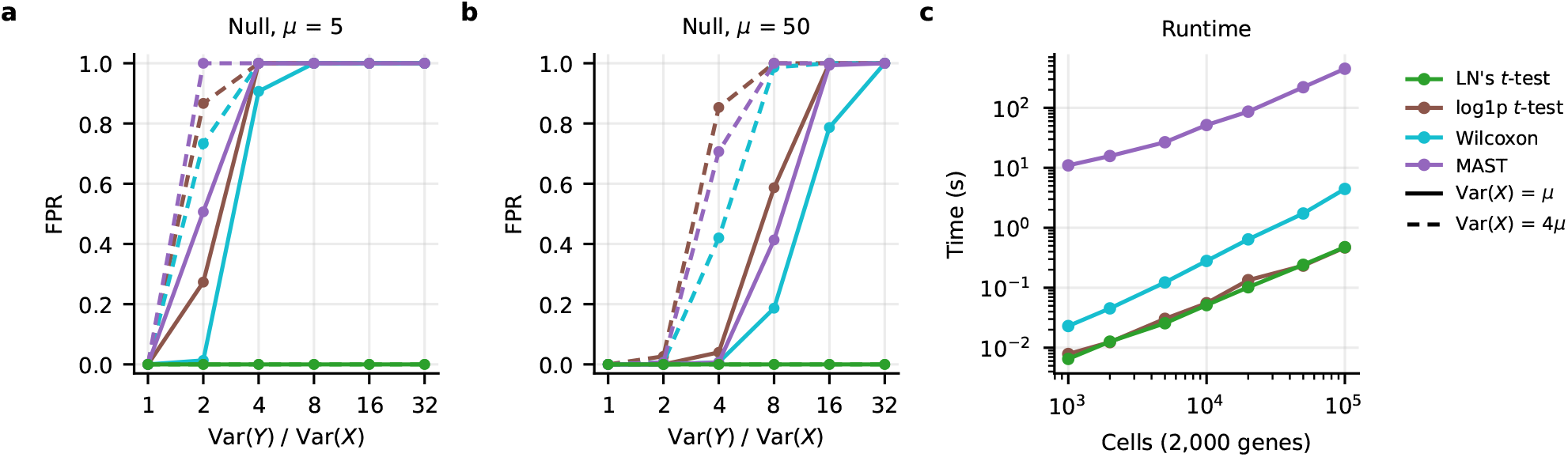
Comparison with MAST and runtime scaling. **a**,**b**, False-positive rate (FPR) of each test against Var(*Y*)*/*Var(*X*) on genes of equal mean, for *µ* = 5 (**a**) and *µ* = 50 (**b**), with Var(*X*) = *µ* (solid) or 4*µ* (dashed). **c**, Median wall time of each test, from raw counts to adjusted *p*-values, against the number of cells.

**Extended Data Figure 3:**
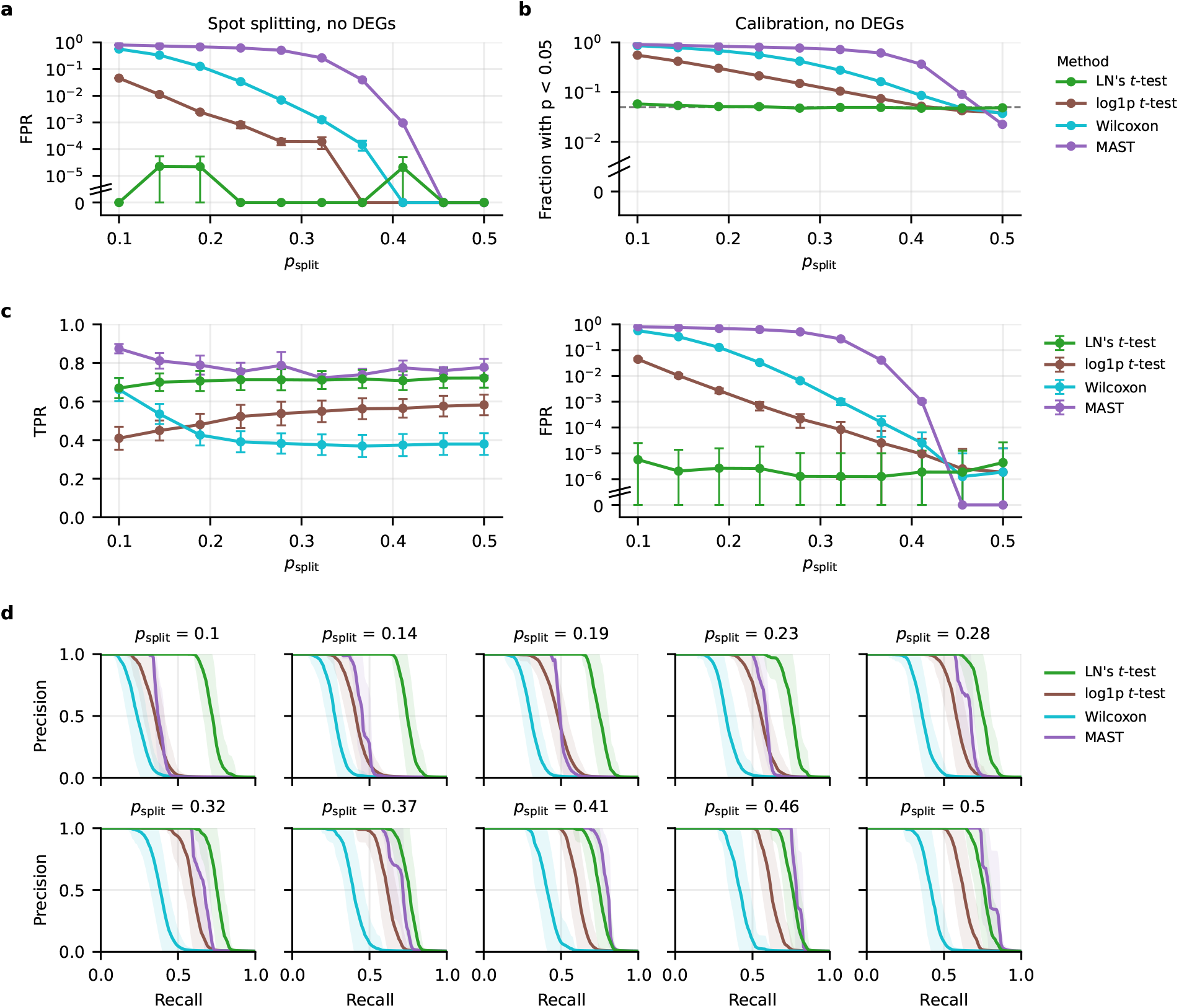
Spot splitting experiment on kidney Visium HD data. **a**, False-positive rate (FPR) without differentially expressed genes (DEGs) as in Figure 1 f, with MAST. **b**, Fraction of genes with uncorrected *p* < 0.05 without DEGs against *p*_split_, where a calibrated test gives 0.05 (dashed). **c**, True-positive rate (TPR, left) and FPR (right) with simulated DEGs, as in Figure 1 g. **d**, Mean precision–recall curves at every *p*_split_. Data are means and standard deviations over replicates.

**Extended Data Figure 4:**
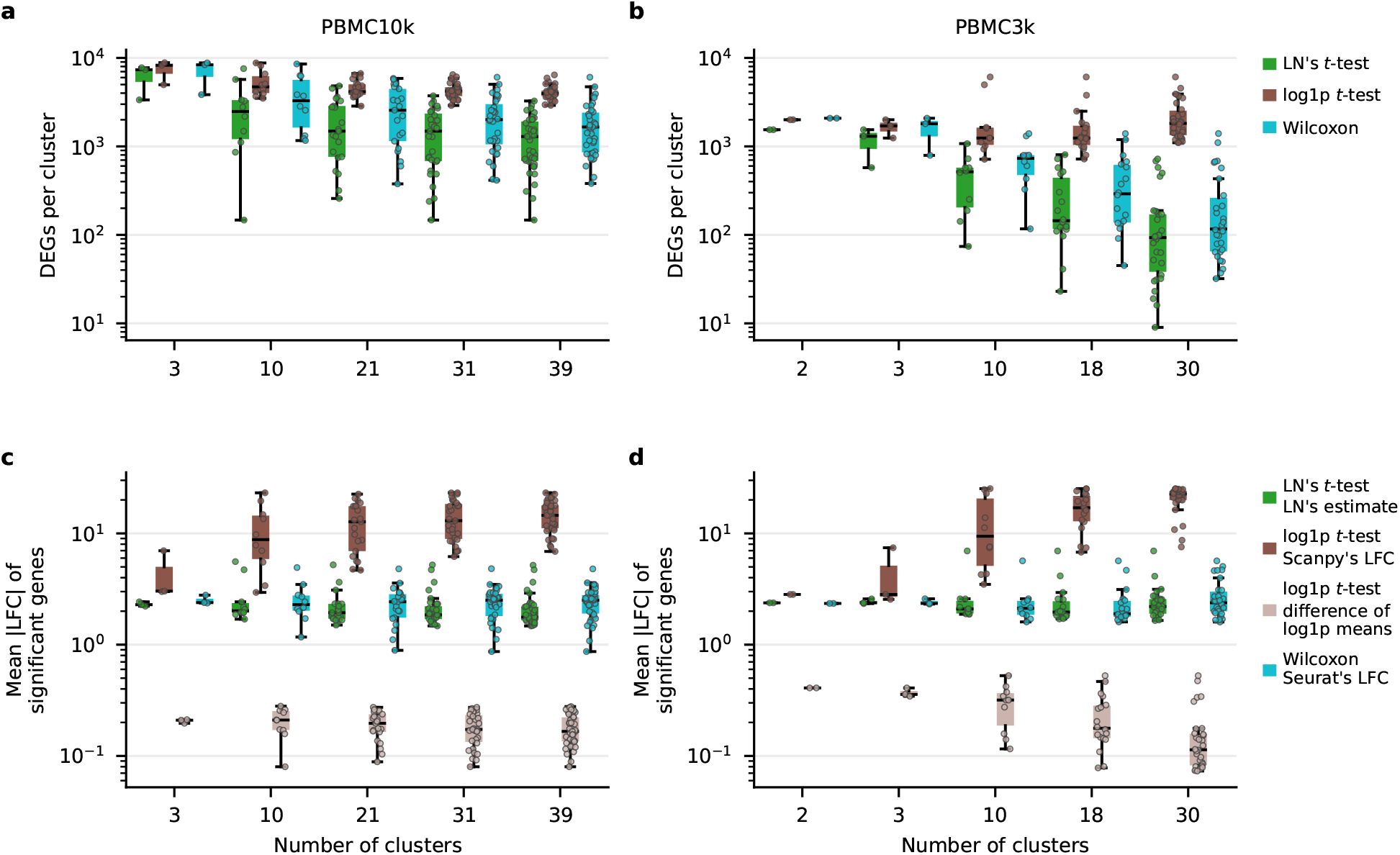
Differentially expressed genes and reported effect sizes of the PBMC clusters. **a**,**b**, Significant genes per cluster for each test, in PBMC10k (**a**) and PBMC3k (**b**). **c**,**d**, Mean absolute log-fold change (|LFC|) of each test’s significant genes per cluster as each package reports it, in PBMC10k (**c**) and PBMC3k (**d**). The lighter boxes show the difference in mean log1p expression that the log1p *t*-test tests. Each point is a cluster, and the boxes summarize the clusters of each resolution, on log axes.

**Extended Data Figure 5:**
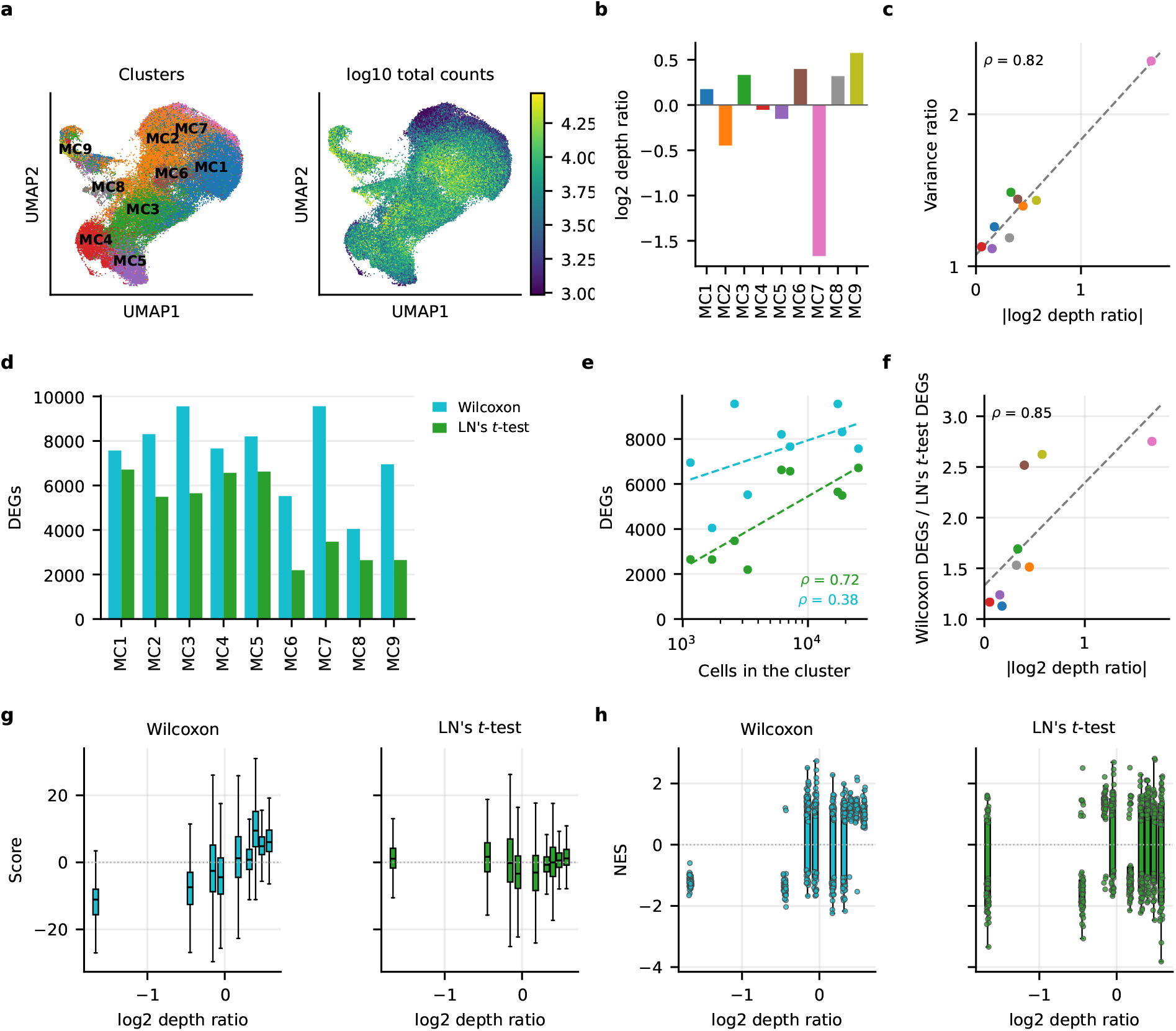
Cluster depth, differentially expressed genes and enrichment in the glioma myeloid cell atlas [42]. **a**, UMAP of the myeloid cells coloured by the clusters of the original study (left) and by log_10_ total counts per cell (right). **b**, Each cluster’s log_2_ depth ratio. **c**, Each cluster’s variance ratio, as in Figure 2 e, against its |log_2_| depth ratio, with a least-squares line (dashed) and the Spearman correlation. **d**, Significant genes per cluster for Wilcoxon and LN’s *t*-test. **e**, Each test’s significant genes against the cluster’s number of cells (log axis), with least-squares lines (dashed) and Spearman correlations. **f**, Ratio of Wilcoxon’s significant genes to LN’s against the log_2_ depth ratio, with a least-squares line (dashed) and the Spearman correlation. **g**, Wilcoxon *z*-scores and LN’s scores of each cluster’s expressed genes against its |log_2_| depth ratio, on one axis. Whiskers extend to 1.5 times the interquartile range, and individual genes are not drawn. **h**, Normalized enrichment scores (NES) of the 50 Hallmark gene sets in each cluster against its log_2_ depth ratio, with genes ranked as in **g** and each gene set a point.

## Supplementary Note A *t*-testing in single-cell transcriptomics

In single-cell transcriptomics, *y* is an *n × g* gene expression matrix, with *n* the number of cells and *g* the number of genes, so that *y*_*ij*_ ∈ [0, ∞) is a unique molecular identifier (UMI), or read count of gene expressions from cell *i* ∈ [1, *n*] and gene *j* ∈ [1, *g*]. It is common practice, however, to not work with the raw counts, but instead normalizing and transforming the data before analysis. A typical combination of normalization technique and transform includes the counts-per-10,000 (CP10K) normalization followed by log1p transform.

In a differential gene expression experiment, one additionally obtains *x* ∈ [0, ∞)^*n′ ×G*^ corresponding to an alternative condition (examples of comparative conditions include treatment vs. control, different cell types, or spatially distinct cell populations), and then asks whether there is a significant LFC between the two conditions with respect to a specific gene *g*. Mathematically, the LFC measures the difference in the log_2_-transformed expected expression values for gene *j* across conditions,

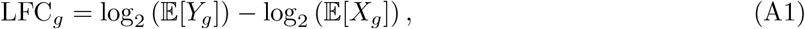

where *Y*_*g*_, *X*_*g*_ are the random variables (r.v.s) that, respectively, follow the condition-specific distributions of the *g*-th gene’s expression values. For ease of exposition, we will drop the gene-specific index going forward.

The expected values are unknown, so they must be approximated using estimators. To decide whether an LFC estimate is significantly different from zero, practitioners employ popular and well-known statistical frameworks, like the *t*-test. In the *t*-test, one constructs a *t* statistic which compares the difference between two inferred means 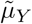 and 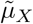 scaled by the standard error of the estimates 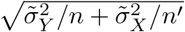, that is

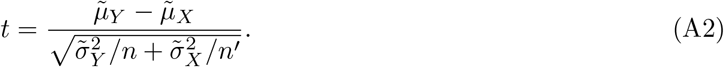

For a critical value, *t*_*α,ν*_, where *α* is a given significance level and *ν* the degrees of freedom, confidence intervals (CIs) that quantify the uncertainty in the inferred difference in means can be obtained following [1]

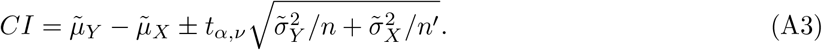

The confidence intervals can also be used to identify differential gene expression (DGE) classification

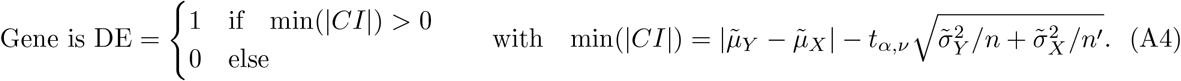

### A.1 *t*-tests on log1p-transformed samples inflate FPRs

Above, we provided a presentation of *t*-testing in single-cell transcriptomics without specifying how the means and variances in Equation (A2) are typically estimated. In both `scanpy` and `Seurat`, these parameters are empirical estimates based on the distribution of the log1p-transformed samples, i.e. 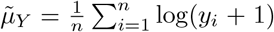 and 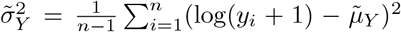, and likewise for *X*. Our first result is to prove that log1p *t*-testing asymptotically produces inflated FPRs under mild assumptions on the gene expression distributions.

#### Theorem 1

*Let* 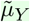 *and* 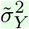 *be unbiased empirical estimates of* E[log(*Y* +1)] *and* Var(log(*Y* +1)), *respectively, and similarly for* 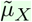 *and* 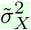, *and assume that Y and X follow negative binomial distributions with finite means and variances such that* E[*Y* ] = E[*X*] *but* Var(*Y*)≠ Var(*X*), *then the corresponding t-test produces type I errors with almost sure convergence in the number of samples, i*.*e*.

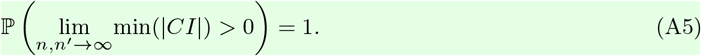

*Proof* See Section C. □

*Remark 1* Theorem 1 trivially extends to pairs of Poisson and negative binomial r.v.s, as the Poisson distribution is a special case of a negative binomial distribution.

*Remark 2* Through [2] (see their Theorem 3 in §3), our Theorem 1 can also be extended to pairs of Beta distributed r.v.s, which may be a suiting distributional assumption on the normalized count data.

*Remark 3* Theorem 1 *does not* extend to pairs of Poisson r.v.s, as the Poisson is a single-parameter distribution. Experimental results predicted by Theorem 1 thus suggest that the associated data is likely not Poisson distributed.

The above sample-asymptotic result is alarming, and suggests that log1p *t*-tests should not be trusted. Another perspective from which we can study this bias is via the difference in variance.

#### Conjecture 1.

*Under the same assumptions as in Theorem 1, FPRs are expected to become inflated for finite sample sizes when the difference in variance increases. That is*, ℙ (min(|*CI*|) > 0) → 1 *as* Var(*X*) *and* Var(*Y*) *become increasingly dissimilar. The following intuition, based on LN r*.*v*.*s, for which closed-form expressions are available, motivates this conjecture*.

Having closed form expressions of the means of the log-transformed samples helps guide the intuition, and so we study the bias using LN r.v.s and the log transform: Let 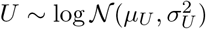 and 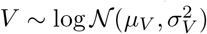, and set E[*U* ] = E[*V* ] = *m* but Var(*V*) < Var(*U*), then 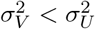 and

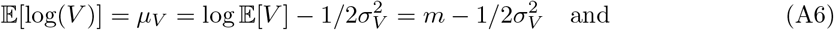

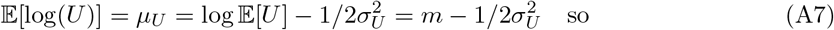

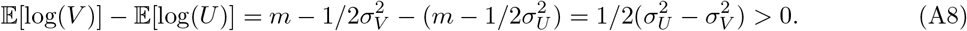

This shows that the gap between the means of the two log-transformed r.v.s increases with the difference in variance. Although this example was constructed using LN distributed r.v.s and the log transform, the intuition carries over to NB r.v.s and the log1p transform since, firstly, pairs LN r.v.s and pairs of NB r.v.s can be concavely ordered under the assumptions in Conjecture 1, while secondly, both log and log1p are concave transforms. Indeed, in our experiments we empirically show that increasing the size of the difference in variance for NB r.v.s correlates to inflated FPRs when using the log1p *t*-test.

*Remark 4* The *t*-tests provided in `scanpy` and `Seurat` are systematically biased in the way described in Theorem 1 and Conjecture 1.

### Supplementary Note B LN’s *t*-test

Here we propose LN’s *t*-test, a differential test based on our new, asymptotically unbiased LFC estimator for which we derive its associated *t* statistic and CIs. The construction of the LFC estimator avoids the log1p-transform, and so LN’s *t*-test does not suffer from inflated FPRs—unlike a *t*-test on log1ptransformed samples.

The proofs of all the lemmas and theorems are available in Section D, and as in Section A we drop the gene index for clarity of exposition.

To construct our estimator, we note that the expected gene expression value can be decomposed as

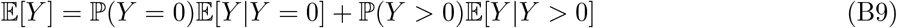

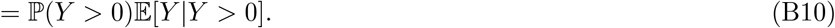

Equation (B10) consists of two terms, the probability of an observation being positive, ℙ(*Y* > 0), and the expected positive gene expression value, E[*Y*|*Y* > 0]. When testing differences in LFC, there are hence four LFC terms to be estimated: ℙ(*Y* > 0), E[*Y*|*Y* > 0] and ℙ (*X* > 0), E[*X* | *X* > 0]. The two subsequent sections are devoted to developing LN distributed estimators of these terms. By doing so, we obtain an LFC estimator in Section B.3 that is normally distributed with unbiasedness guarantees, and closed-form confidence intervals.

### B.1 The Estimator of *P* (*Y* > 0)

Supported by the Bayesian hypothesis testing literature (§3.10.2.4 in [3]), we make the assumption that *P* (*Y* > 0) can be accurately estimated by a Beta distributed r.v.,

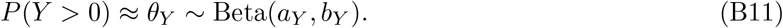

This is a sensible choice, not least due to its constrained range. However, recall that we request our estimator of *P* (*Y* > 0) to be LN distributed.

To this end, we now show a novel LN approximation of the Beta probability density function (pdf) by noting that the mean and variance of a log-transformed Beta r.v. enjoy closed-form expressions,

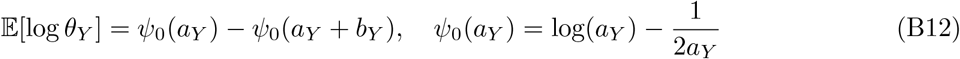

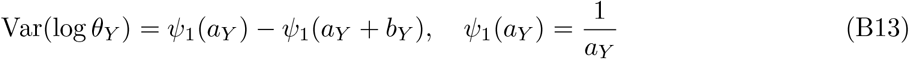

where *ψ*_0_ and *ψ*_1_ are matched, truncated large-argument asymptotic expansions of the digamma and trigamma functions, respectively [4, Eq. 5.11.2 and 5.15.8]. Using the expressions, we approximate the Beta(*a*_*Y*_ , *b*_*Y*_) pdf by an LN pdf 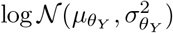 with parameters,

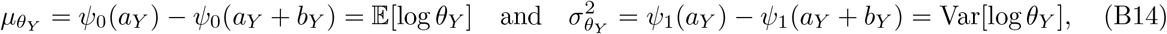

We find that this is a remarkably accurate approximation, especially when the Beta distribution has non-negative skewness (i.e., when *a*_*Y*_ ≤ *b*_*Y*_ ; see Supplementary Figure 1 for an illustrative example), and so, from here, we assume that *θ*_*Y*_ is LN distributed. Consequently, the mean of *θ*_*Y*_ is

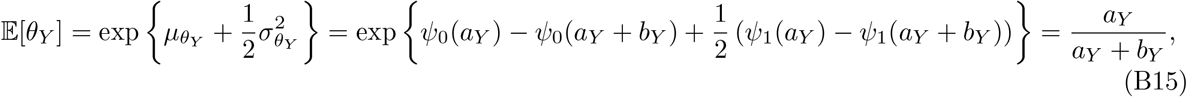

which, notably, is also the mean of the Beta(*a*_*Y*_ , *b*_*Y*_) pdf.

Unfortunately, *a*_*Y*_ and *b*_*Y*_ are unknown, but, since they are Beta parameters, they can be estimated in an unbiased manner with data, *y*,

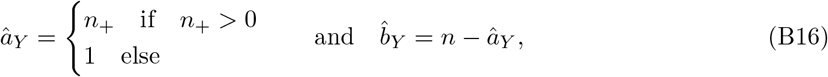

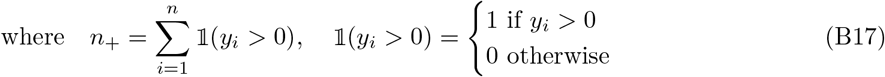

Finally, we choose 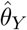4 such that it approximates the LN mean (Equation (B15)), that is we choose

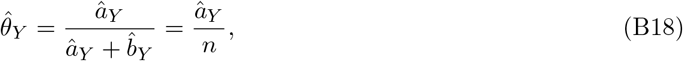

which ensures that it is an asymptotically unbiased estimator of *P* (*Y* > 0).

#### Lemma 1.

*Assuming that P* (*Y* > 0) > 0, 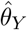 *is an asymptotically unbiased estimator of P* (*Y* > 0).

### B.2 The Estimator of E[*Y* |*Y* > 0]

In accordance with Equation (B17), we refer to the number of samples with positive counts as *n*_+_ and term the index set of these samples as 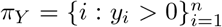. Here we have filtered out zero-counts, so we can log-transform the sample mean of the remaining, positive samples without adding any pseudocount. This sample mean is consequently strictly positive, and so, motivated by models like the Poisson LN in scRNA-seq (see, e.g., [5] where the mean of the Poisson expression distribution is log-normally distributed), and the log-normal Cox process [6], we let 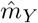 denote the sample mean and assume that it is LN distributed, i.e.

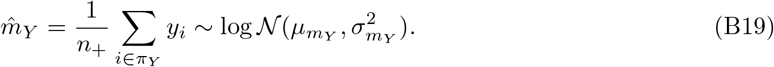

We emphasize that modeling 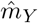 as LN distributed does not imply an LN assumption on the individual samples, *y*_*i*_. In fact, a sum of LN random variables, scaled or otherwise, is not LN in general and so Equation (B19) suggests that the positive gene expressions are *not* LN distributed.

We parameterize the distribution in Equation (B19) through a standard procedure using unbiased empirical estimates of the mean and variance of the LN data (see Section F.1 for a derivation). As a result, 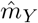 follows an LN distribution with estimated parameters

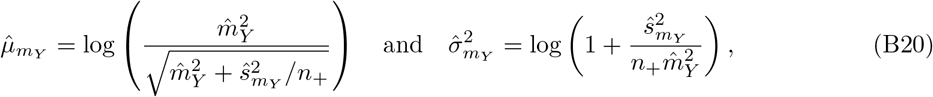

and is clearly unbiased since 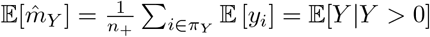.

### B.3 Our LFC estimator is normally distributed

By estimating each of the terms using LN distributed estimators, we can leverage the following lemma.

#### Lemma 2.

*If* 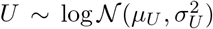 *and* 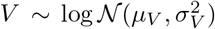 *are independent, then* 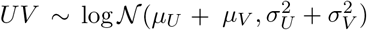 *and* 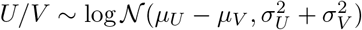.

From the lemma it follows that, if *U, V, U*^*′*^, *V* ^*′*^ (i.e. our estimators of *P* (*Y* > 0), E[*Y* |*Y* > 0], *P* (*X* > 0), E[*X*|*X* > 0]) are independent LN distributed r.v.s, then log_2_(*UV*) − log_2_(*U*^*′*^*V* ^*′*^), is normally distributed.

That is, our proposed LFC estimator follows a normal distribution

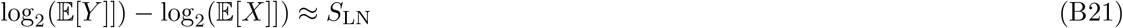

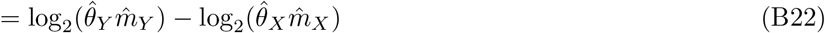

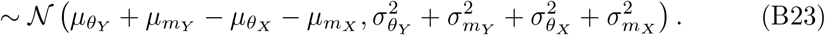

We will now establish that our LFC estimator converges to the true LFC in probability, and that it is asymptotically unbiased. Since Equation (A1) is undefined if P(*Y* > 0) = 0 or P(*X* > 0) = 0, let us safely assume that P(*Y* > 0) > 0 and P(*X* > 0) > 0.

#### Lemma 3.

*Assume that* ℙ(*Y* > 0) > 0 *and* ℙ(*X* > 0) > 0, *then as n and n*^*′*^ *tend to infinity, S*_*LN*_ *converges in probability to Equation* (A1).

Consequently, *S*_LN_ is a consistent estimator of the LFC. By adding mild conditions regarding bounded second moments, we can furthermore state that *S*_LN_ is asymptotically unbiased.

#### Theorem 2

(**Asymptotic Unbiasedness**) *Assume that* ℙ(*Y* > 0) > 0 *and* ℙ(*X* > 0) > 0, *and that* E[*Y* ^2^] < ∞ *and* E[*X*^2^] < ∞. *Then S*_LN_ *is an asymptotically unbiased estimator of Equation* (A1), *i*.*e*.

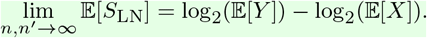

In contrary to the bias of the log1p *t*-test formalized in Theorem 1, Theorem 2 above implies that as we observe more cells, we can be sure that the our LFC estimator and its corresponding confidence intervals (defined in the subsequent section) are centered around the true LFC.

### B.4 Deriving LN’s *t*-test

Given the standard errors

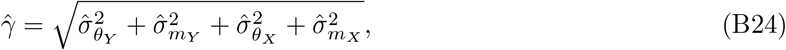

and a practitioner-chosen critical value *t*_*α,ν*_, we can now naturally construct CIs since the LFC estimator in Equation (B21) is normally distributed

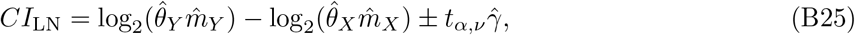

and a *t* statistic

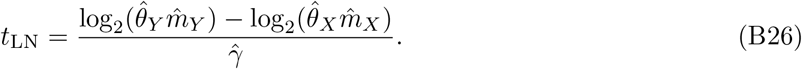

We define LN’s *t*-test as a DGE test based on *CI*_LN_ or *t*_LN_. Algorithm 1 constitutes a pseudo-code pipeline for implementing LN’s *t*-test, showcasing its ease of implementation (see Section F.2 for Algorithm 2 and a time-complexity analysis).

#### Algorithm 1

**LN’s** *t***-test** (gene wise)

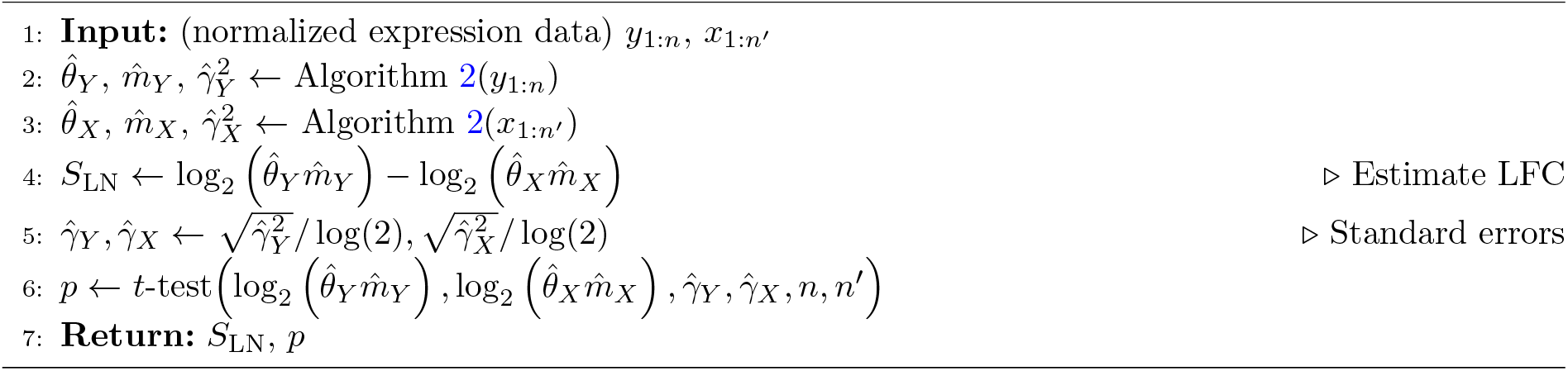

#### Algorithm 2

**Get Estimators**

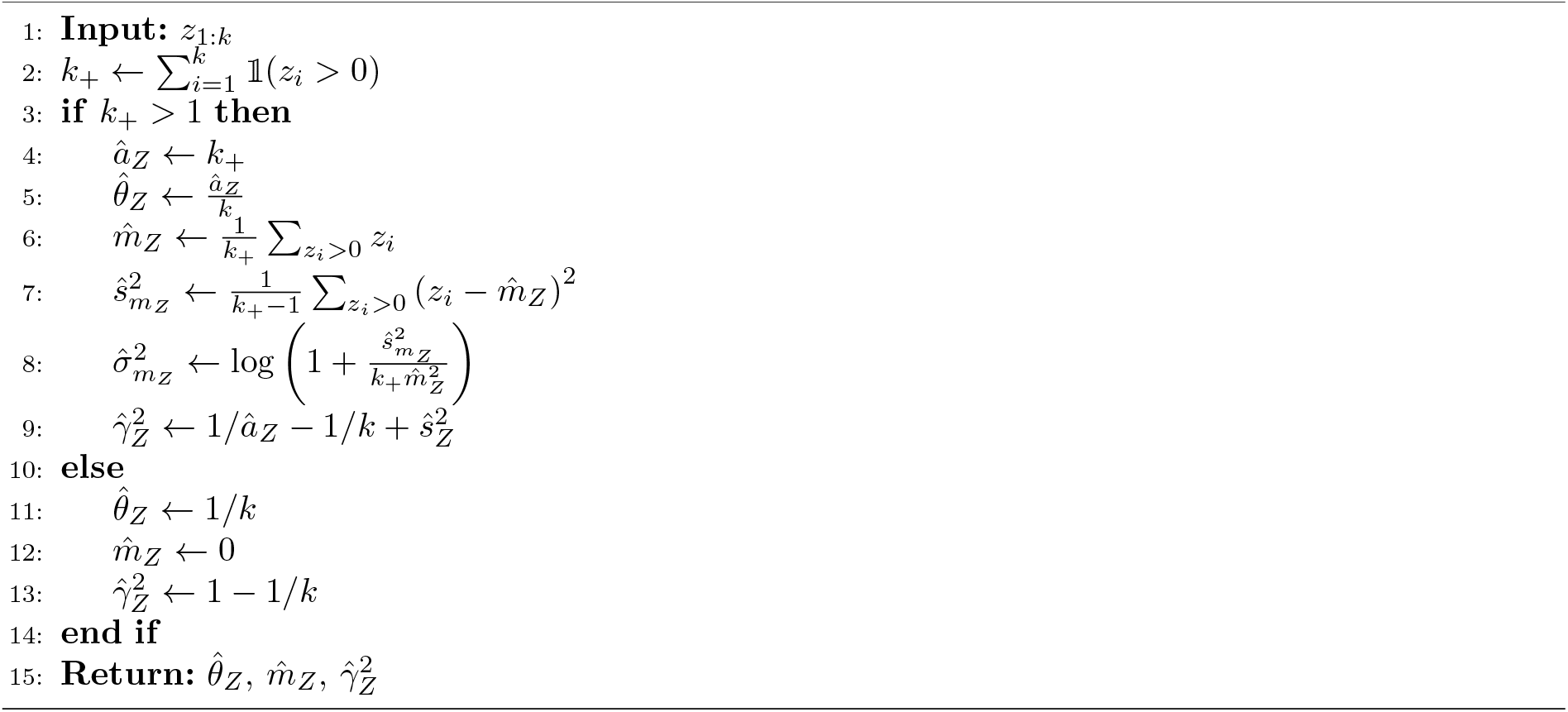

## Supplementary Note C Lemmas and proof of Theorem 1

Lemma 4 and Lemma 5 are notationally rewritten versions of two theorems in [7], while Lemma 6 and Lemma 7, on the other hand, are new results as far as we are aware. We use these lemmas in order to arrive at our new, main theoretical result in Theorem 1.

***Proof sketch***

Lemma 4 provides us with a tool that allows us to leverage Jensen’s inequality, relating the means of log1p-transformed r.v.s — as long as one of the r.v.s is a mean-preserving spread of the other. This result is important since it allows us to prove Theorem 1: In the limit of *n, n*^*′*^ evaluated in Equation (A5), we only need to compare the absolute difference in means of log1p-transformed r.v.s.

So, in order to utilize the martingale property in Lemma 4, we use Lemma 5 from [7] and devise Lemma 6 to show that, under relevant assumptions on two negative binomial r.v.s *X* and *Y* , *Y* is a meanpreserving spread of *X*, or vice versa. We are then ready to relate the means of the log1p-transformed *X* and *Y* by an inequality in Lemma 7, a key step towards proving our main result in Theorem 1.

### Lemma 4

(Theorem 3.A.4 in [7]). *Y* ∼ *p*_*Y*_ *is a mean-preserving spread of X* ∼ *p*_*X*_ *if and only if there exists a joint distribution p*_*X,Y*_ *with marginal distributions such that X*^*′*^ ∼ *p*_*X*_ *and Y* ^*′*^ ∼ *p*_*Y*_ , *that is*, 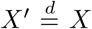 *and* 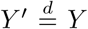 *(equal in distribution), and* {*X*^*′*^, *Y* ^*′*^} *is a martingale, i*.*e*. E[*Y* ^*′*^|*X*^*′*^] = *X*^*′*^ *almost surely*.

### Lemma 5

(Condition 3.A.57 in Theorem 3.A.44 in [7]). *Let X and Y be two random variables with* E[*X*] = E[*Y* ], *and probability mass functions p*_*X*_ *and p*_*Y*_ , *and let a*(*k*) = *p*_*Y*_ (*k*) − *p*_*X*_ (*k*) *be a real-valued function while S*(*a*(*k*)) *denotes the number of sign changes in a*(*k*) *over k, ignoring zero terms. Then Y is a mean-preserving spread of X if S*(*a*(*k*)) = 2 *and the sequence of sign changes is* (+, −, +).

### Lemma 6.

*Let X and Y be negative binomial random variables with the same mean* 0 < *µ* < ∞ *and different dispersions* 0 < *ϕ*_*X*_ < *ϕ*_*Y*_ < ∞ *under the mean–dispersion parameterization*

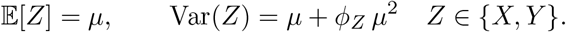

*Then Y is a mean-preserving spread of X*.

*Proof* Let *p*_*Z*_ (*k*) be a negative binomial pmf and *R*(*k*) = *p*_*Y*_ (*k*)*/p*_*X*_ (*k*) for *k* ≥ 0. The negative binomial recurrence gives

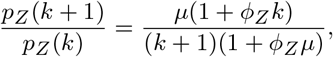

and hence

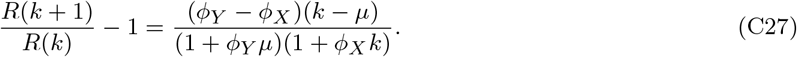

Thus, if *µ* is non-integer, *R*(*k*) has a single minimum as it decreases while *k* < *µ* and increases while *k* > *µ*. If *µ* is integer, then *R*(*µ*) = *R*(*µ* + 1). Consequently, the *R* values below 1 form one consecutive block (irrespective of *µ* being integer or not).

At the left endpoint,

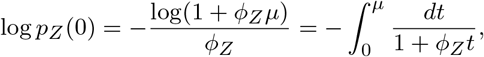

so *p*_*Y*_ (0) > *p*_*X*_ (0) and *R*(0) > 1. At the other end,

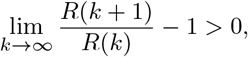

which implies *R*(*k*) → ∞: eventually each successive ratio exceeds some fixed number greater than 1.

Finally, *R*(*k*) < 1 for at least one *k*, otherwise the pmfs *p*_*Y*_ (*k*) and *p*_*X*_ (*k*) cannot both sum to 1. Recalling that *R*(*k*) has a single minimum, *p*_*Y*_ (*k*) − *p*_*X*_ (*k*) therefore has exactly the sign pattern (+, −, +). Since the means are equal, Lemma 5 implies that *Y* is a mean-preserving spread of *X*. □

### Lemma 7.

*Let X and Y be negative binomial random variables with the same mean* 0 < *µ* < ∞ *and different dispersions* 0 < *ϕ*_*X*_ < *ϕ*_*Y*_ < ∞ *under the mean-dispersion parameterization (i*.*e*., *same as in Lemma 6. Furthermore, let φ be any concave function (e*.*g*., *φ*(*z*) = log(*z* + 1)*), then* E[*φ*(*Y*)] < E[*φ*(*X*)].

*Proof* From Lemma 6 we know that *Y* is a mean-preserving spread of *X*, and so we can combine the martingale property from Lemma 4 with Jensen’s inequality to show that

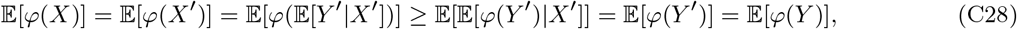

where strict inequality is achieved in any non-degenerate case, i.e. whenever Var(*Y*) > Var(*X*) which is the case here.

Exemplifying with the special case of *φ* being the log1p transform, we have thus shown that

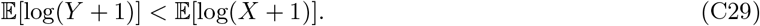

□

### Theorem 1

*Let* 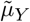 *and* 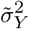 *be unbiased empirical estimates of* E[log(*Y* + 1)] *and* Var(log(*Y* + 1)), *respectively, and similarly for* 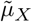 *and* 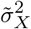, *and assume that Y and X follow negative binomial distributions with finite means and variances such that* E[*Y* ] = E[*X*] *but* Var(*Y*) ≠ Var(*X*), *then the corresponding t-test produces type I errors with almost sure convergence in the number of samples, i*.*e*.

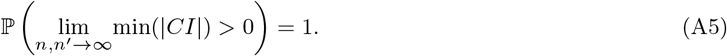

*Proof* Using the notation from Equation (A4), we need to prove Equation (A5), in other words, that the probability of falsely calling a gene differentially expressed converges to one with the number of samples. We will rely on the law of large numbers, and Lemma 7.

According to the strong law of large numbers 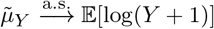, and vice versa for 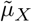, while

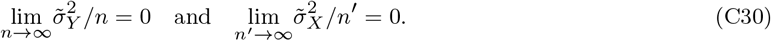

Hence, we need to show that the following inequality holds

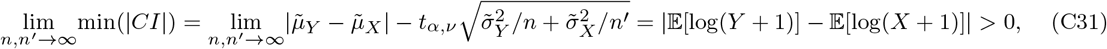

which follows from Lemma 7 as all the conditions are met: if Var(*X*) > Var(*Y*) then,

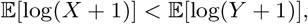

and if Var(*X*) < Var(*Y*), then instead

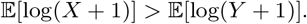

Hence the following has been shown

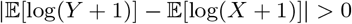

which, under the assumptions on *X* and *Y* , proves that

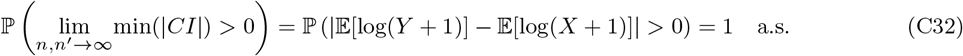

□

## Supplementary Note D Proofs of statements in the derivation of LN’s *t*-test

In this section Lemma 1, Lemma 2, Lemma 3 and Theorem 2 are restated and proved.

### Lemma 1.

*Assuming that P* (*Y* > 0) > 0, 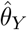 *is an asymptotically unbiased estimator of P* (*Y* > 0).

*Proof* The assumption *P* (*Y* > 0) > 0 implies that *P* (lim_*n*→∞_ *n*_+_ = 0) = 0, and so, by the law of large numbers, 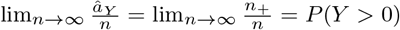. □

### Lemma 2.

*If* 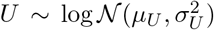 *and* 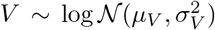 *are independent, then* 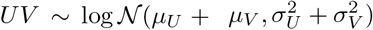 *and* 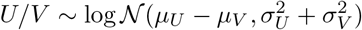.

*Proof* The log of *UV* is a sum of two normally distributed r.v.s,

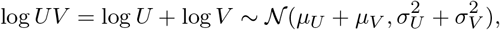

and so

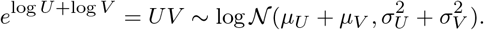

This extends trivially to *U/V*

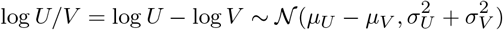

yielding

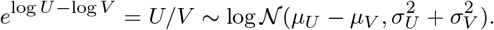

□

### Lemma 3.

*Proof* Since (law of large numbers) 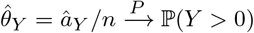 and 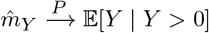, we have 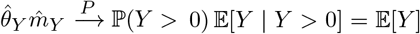, and similarly for *X*. Thus, by continuity of log_2_, 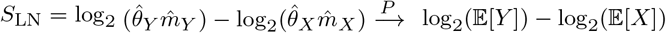. □

**Theorem 2** (**Asymptotic Unbiasedness**) *Assume that* ℙ(*Y* > 0) > 0 *and* ℙ(*X* > 0) > 0, *and that* E[*Y* ^2^] < ∞ *and* E[*X*^2^] < ∞. *Then S*_LN_ *is an asymptotically unbiased estimator of Equation* (A1), *i*.*e*.

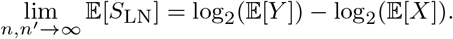

*Proof* By Lemma 3, 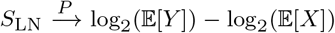. Since E[*Y* ^2^] < ∞ (and likewise for *X*), the sample mean 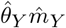 converges to E[*Y* ] in *L*_1_ norm (a.k.a. convergence in mean): Using Cauchy-Schwarz to upper-bound the mean absolute error,

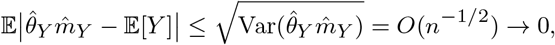

and the same holds for 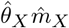. The function log_2_ is continuous, so

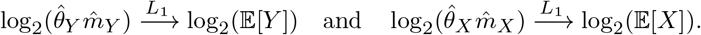

Meanwhile, convergence in *L*_1_ implies convergence of expectations, i.e.

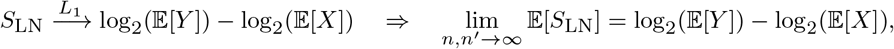

which concludes the proof. □

## Supplementary Note E The log-normal approximation of the Beta pdf for large *P* (*X* > 0)

The support of the log-normal distribution is not upper-bounded, and so we investigate the behavior of our new approximation of the Beta pdf when *P* (*X* > 0) is closer to one, i.e. when *a* > *b*. Notably, the mean of our estimator is the maximum likelihood estimator of the Beta mean. As such, our estimator is consistent, and unbiased (as shown above). Furthermore, the contribution from the estimator to the standard error in Equation (B24) rapidly goes to zero as *a* increases, even with *b* = 1 (i.e, a single zero count; see Supplementary Figure 2). For *b* > 1 the standard errors decay at a higher rate. As such, we do expect LN’s *t*-test to be robust in high non-zero count regimes.

Out of interest, and to provide a more holistic analysis of our moment-matched log-normal pdf, we also analyze the Kullback-Leibler (KL) divergence between the Beta pdf, and a log-normal pdf with our specific parameterization.

Let *q*(*θ*) = Beta(*θ* | *a, b*) on *θ* ∈ (0, 1), i.e.

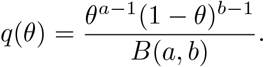

Let *p*(*θ*) = log *N* (*θ* | *µ, σ*^2^) with

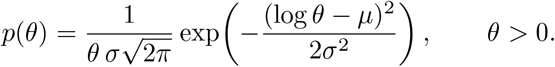

**Supplementary Figure 1:**
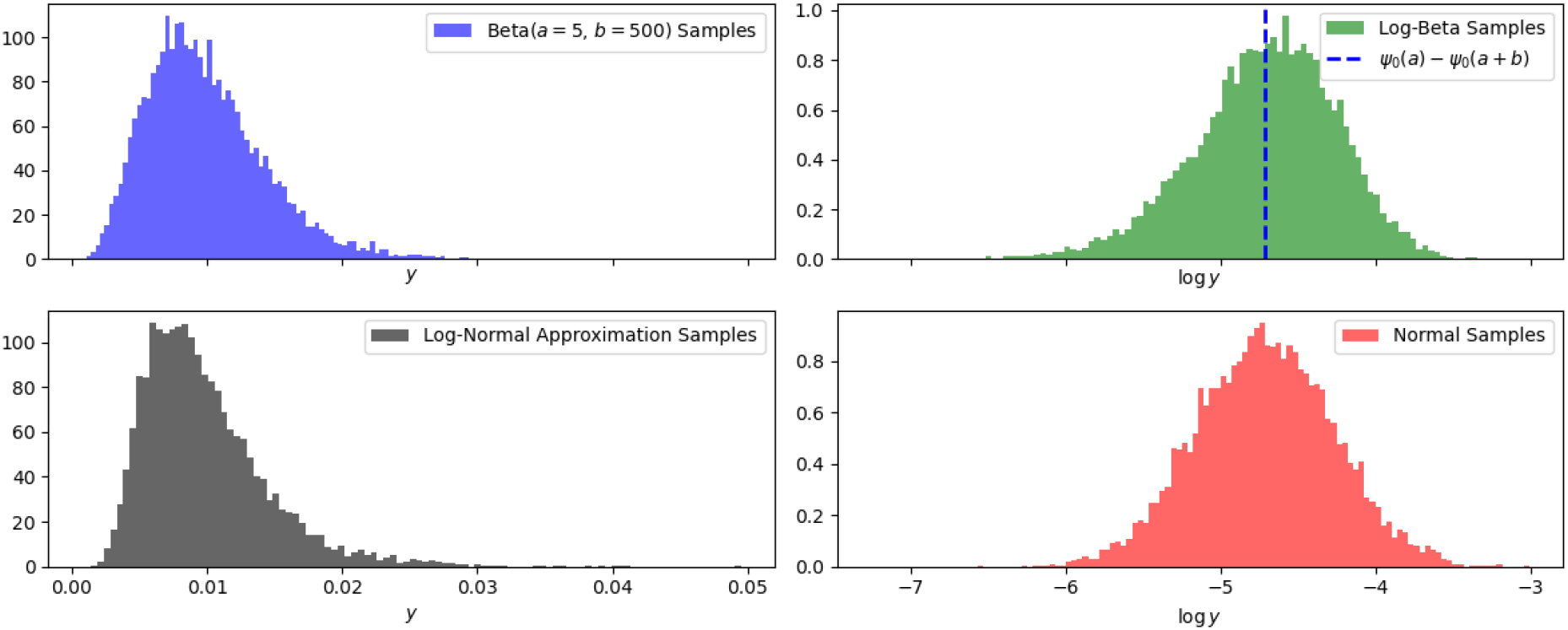
Visualization of the similarities of a Beta distribution and its log-normal approximation. The bottom axes are aligned in each column. **Top left:** Samples drawn from a Beta(*a* = 5, *b* = 500). **Top right:** The log-transformed samples from the Beta distribution and the estimated mean of the distribution (dashed blue). **Bottom left:** Samples drawn from log *N* (*ψ*_0_(*a*) − *ψ*_0_(*a* + *b*), *ψ*_1_(*a*) − *ψ*_1_(*a* + *b*). **Bottom right:** Samples drawn from *N* (*ψ*_0_(*a*) − *ψ*_0_(*a* + *b*), *ψ*_1_(*a*) − *ψ*_1_(*a* + *b*).

**Supplementary Figure 2:**
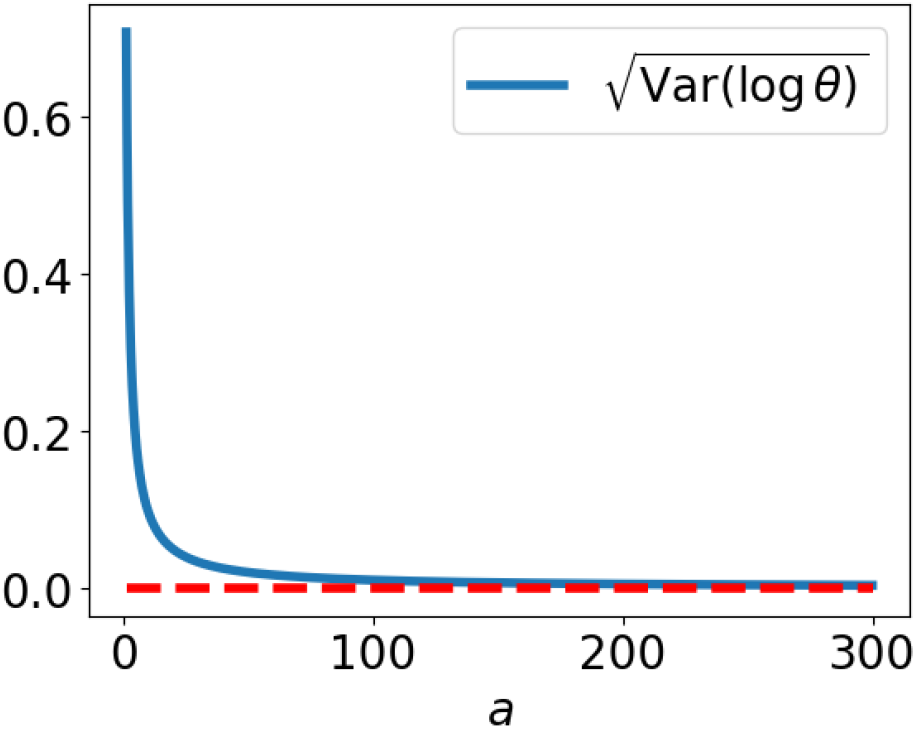
Keeping *b* = 1 fixed and increasing *a*, the standard errors of the *P* (*Y* > 0) estimator rapidly decrease to zero (red, dashed line), as requested. For *b* > 1 the standard errors decay at a higher rate.

Then

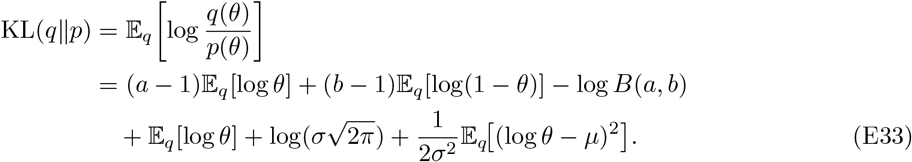

**Supplementary Figure 3:**
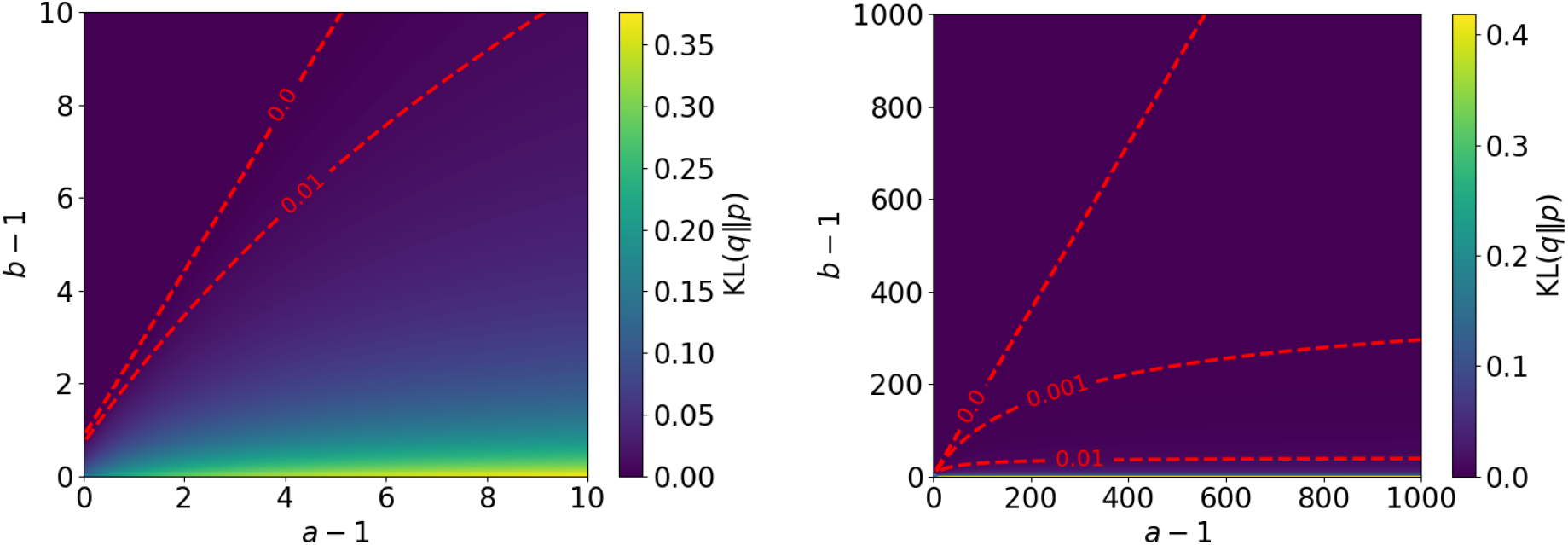
A 2D KL(*q p*) map for different values of (*a, b*) (see Equation (E34).) The dashed red lines mark the boundary where the KL is zero and less than 0.01 or 0.001. The right plot is a zoomed-out version of the left plot.

Using the standard Beta identities

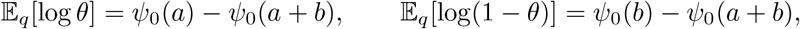

and

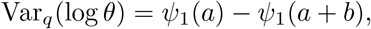

we have

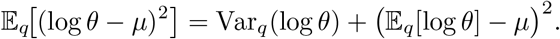

In particular, for our LN approximation (moment-matching on log *θ*),

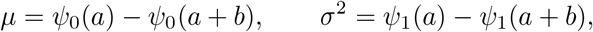

so E_*q*_[(log *θ* − *µ*)^2^] = Var_*q*_(log *θ*) = *σ*^2^, and (E33) simplifies to

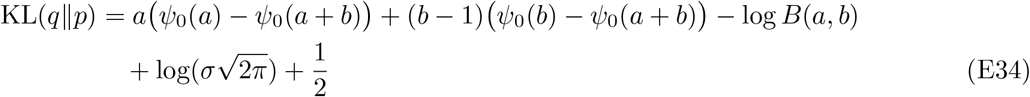

In Supplementary Figure 3 we plot the 2D KL map for different values of (*a, b*). As expected, the KL is largest in the regime where *b* is small and *a* > *b*. However, the KL rapidly decreases with *b*, and is less than 0.01 for *b* > 40 and any *a* (although not shown here, we pushed *a* up to 10,000 without crossing this threshold). Remarkably, there are *b* > *a* regions (corresponding to sparse expression data regimes in single-cell transcriptomics terms) where the KL is numerically zero. Analytically defining this decision boundary is left as future work, but the boundary’s existence highlights the significance of LN’s *t*-test from a broader statistics perspective.

Less rigidly, we finally plot Beta, log-Beta, log-normal and normal samples in Supplementary Figure 1 for a single parameter setup to visualize the resemblance of the distributions.

## Supplementary Note F LN’s *t*-Test and Our Proposed LFC Estimator

Here we provide some additional information regarding the derivations in Section B.

### F.1 The system of equations

This is the full formulation of the system of equations solved in Section B.2.

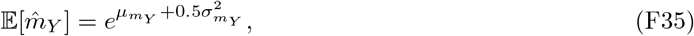

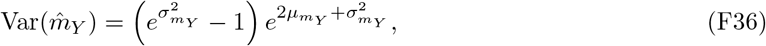

resulting in

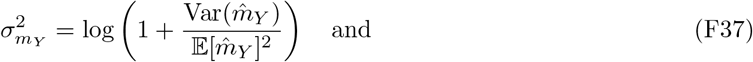

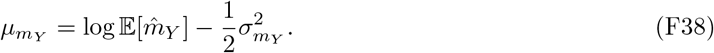

Substituting the unknown terms with our approximations, i.e. 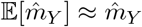 and

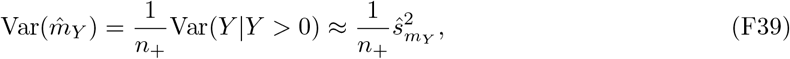

where

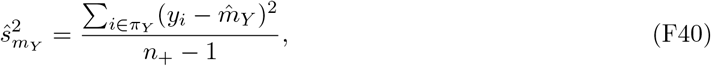

we can write

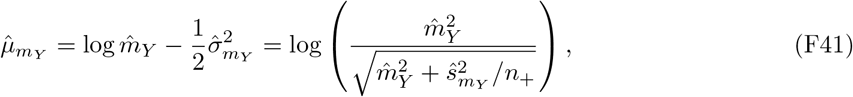

and

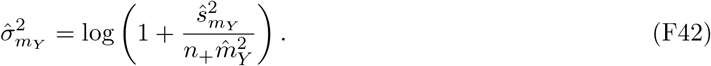

### F.2 Time complexity

Algorithm 1 and 2 consist of series of *O*(1), *O*(*n*), *O*(*n*_+_), *O* (*n*^*′*^) and 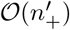 operations. Since *O*(*n*) ≥ *O*(*n*_+_) and 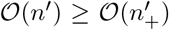, summing the costs of these operations results in an *O* (*n* + *n*^*′*^) complexity. Algorithm 1 is called for all *G* genes so the total complexity of LN’s *t*-test is (*G*(*n* + *n*^*′*^)), which should be the same for the *t*-test implementations in scanpy and seurat.

## Supplementary Note G A digamma-based intuition for small log-scale effect sizes and tiny standard errors

After depth normalization (e.g. CP10K), each cell is rescaled by its library size so that the total expression across genes is approximately constant. Consequently, the per-gene normalized expression can be viewed as a *proportion* of the cell’s total transcriptome. A convenient idealization is therefore to treat the vector of normalized expressions as Dirichlet-distributed, and each gene marginally as approximately Beta-distributed. This approximation is not meant as a generative model of raw counts, but as an analytically tractable proxy for the distribution of *normalized* non-negative values in the unit interval.

To keep the derivation transparent, we use the log transform rather than log 1*p*: the same qualitative conclusions follow from concavity for log 1*p*, but log admits closed-form identities under Beta/Dirichlet models (digamma and trigamma), which makes the bias–variance interplay explicit.

Let *X*_*g*_ ∼ Beta(*a*_*X*_ , *b*_*X*_) and *Y*_*g*_ ∼ Beta(*a*_*Y*_ , *b*_*Y*_) denote depth-normalized expression for gene *g* in the two groups. Define the (natural-log) effect size

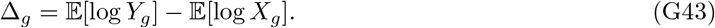

Under the Beta model,

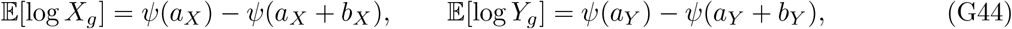

where *ψ* is the digamma function. In contrast, LN’s target estimand is the log-fold change between *population means*

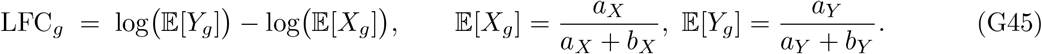

Using the matched, truncated large-argument asymptotic expansions of the digamma and trigamma functions introduced in Section 6.2 (i.e. 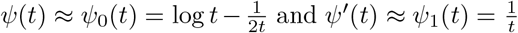; where we denote *ψ*^*′*^(*t*) as the trigamma function), we obtain the approximation

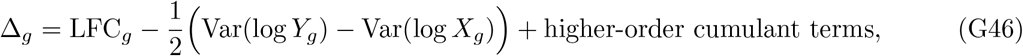

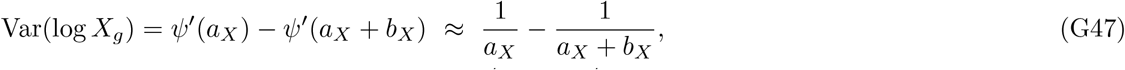

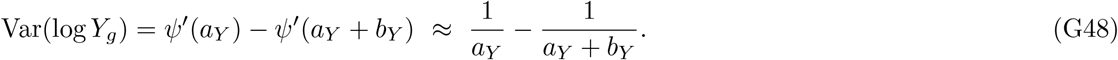

### G.1 Why Δ_***g***_ **can be small even when LFC**_***g***_ ≠ **0**

Equation (G46) makes explicit that the log-scale effect size Δ_*g*_ is not only a function of the LFC, but also of the *difference in log-scale variances* between groups. Hence, even when LFC_*g*_ ≠ 0, a compensating difference in Var(log *·*) can shrink Δ_*g*_ toward zero. In fact, clearly there are cases where Δ_*g*_ = 0 although LFC_*g*_ ≠ 0, which additionally means that unequal variance may cause the log1p *t*-test to produce false *negatives*.

### G.2 The bias is a bug, not a feature

Although a small effect size might seem favorable when measuring the expression difference between samples from a homogeneous population, the example above shows that the effect size of the log1p *t*-test is (approximately) a linear combination of the LFC term and the difference in the variance of the log-transformed samples. As such, the magnitude of the effect size is confounded, leaving the source of a log1p *t*-test result challenging to analyze.

Moreover, we observe small standard errors in our experiments, which explains the stunning number of false discoveries from the log1p *t*-test, despite the small effect sizes.

## Supplementary Note H The high false-positive rates of the Wilcoxon rank-sum test are expected

In formal terms, to see why the Wilcoxon rank-sum test calls a gene DE when the two populations have the same mean but different variances, we first describe what the test is measuring. Let *X* and *Y* denote r.v.s from the two separate gene-expression populations, then the Wilcoxon rank-sum test uses a consistent estimator of *ϑ*, where

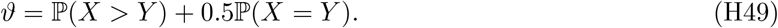

If the estimator of *ϑ* is significantly different from 0.5, then the test calls the considered gene DE [8].

Although *ϑ* cannot be analytically computed for pairs of NB distributed r.v.s, we can understand the degenerating Wilcoxon rank-sum test results observed in our experiments through the conjecture below, which is enabled via LN assumptions on the populations.

An LN assumption on gene expression distributions is a strong assumption when the data contains zero counts. However, our aim here is not to provide a rigid theoretical result, but a statistical intuition of why the false positive rates tend to very high numbers under the considered conditions. For this, the LN assumption is sensible because of the mean-variance dependencies in both NB and LN distributions, their non-negative supports, and because the LN property in Lemma 2 gives an analytical solution to *ϑ*.

### Proposition 3

*Let* 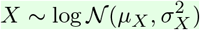 *and* 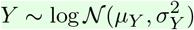 *with* E[*X*] = E[*Y* ], *then if* 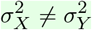, *ϑ* = ℙ(*X* > *Y*) + 0.5ℙ(*X* = *Y*) ≠ 0.5.

*Proof* First, note that if 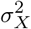 and 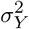 are different, then there is a difference in variance as E[*X*] = E[*Y* ], according to the assumptions. Let *Z* = *X/Y* and recall that via Lemma 2 we have

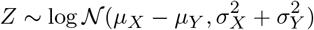

where

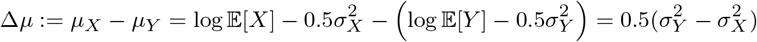

which is non-zero for different variances. From here on, without loss of generality, assume that 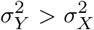. Then, via the monotonicity of the log-transform, and since *X* and *Y* are continuous r.v.s, we conclude that

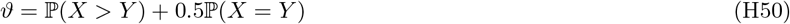

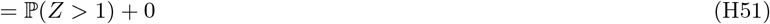

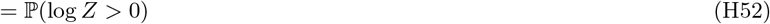

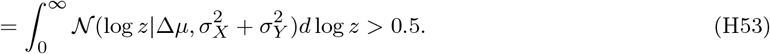

□

## Supplementary Note I The LFC estimator in scanpy

In a recent work [9], an expression for the batch-effect adjusted version of the LFC estimator in `scanpy` was provided as

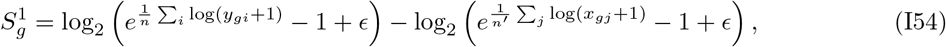

with *ϵ* = 10^−9^. Notably, *ϵ* is added to avoid log_2_(0) but its effect does not diminish with the number of cells. Instead, we can observe massive log-fold change estimates based on 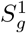 when the data is sparse, attributed to the log_2_(*ϵ*) bias.

This quantity is the `logfoldchanges` value associated to a gene returned from the `rank_genes_groups` function. Note that this is not the same value as the effect size used to test significance in the log1p *t*-test.

## Supplementary Note J Supplementary Results

### Supplementary Results 1: Kidney

These experiments add two negative controls to the spot splitting of Figure 1 e–h, on the same podocyte-containing glomerular capsules: capsules split by sequencing depth alone, and the same split after downsampling every capsule’s unique molecular identifiers (UMIs). As the capsules are biologically homogeneous, every differentially expressed gene (DEG) called without simulated DEGs is a false positive.

**Supplementary Figure 4:**
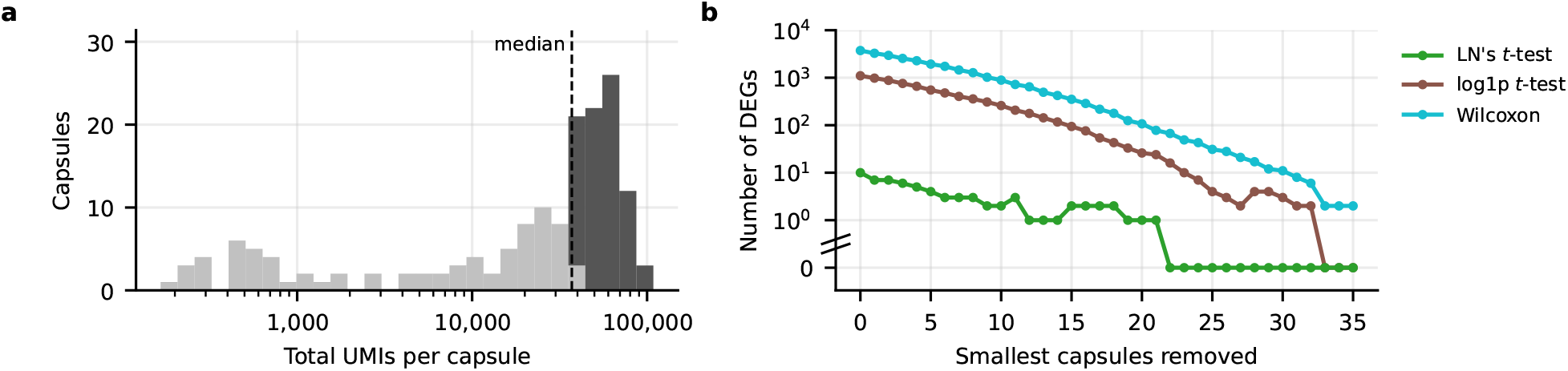
Kidney capsules split by sequencing depth. **a**, Total UMIs per capsule, split at the median into a shallower and a deeper half. **b**, DEGs each test calls between the deeper and the shallower half, as the smallest capsules are removed.

**Supplementary Figure 5:**
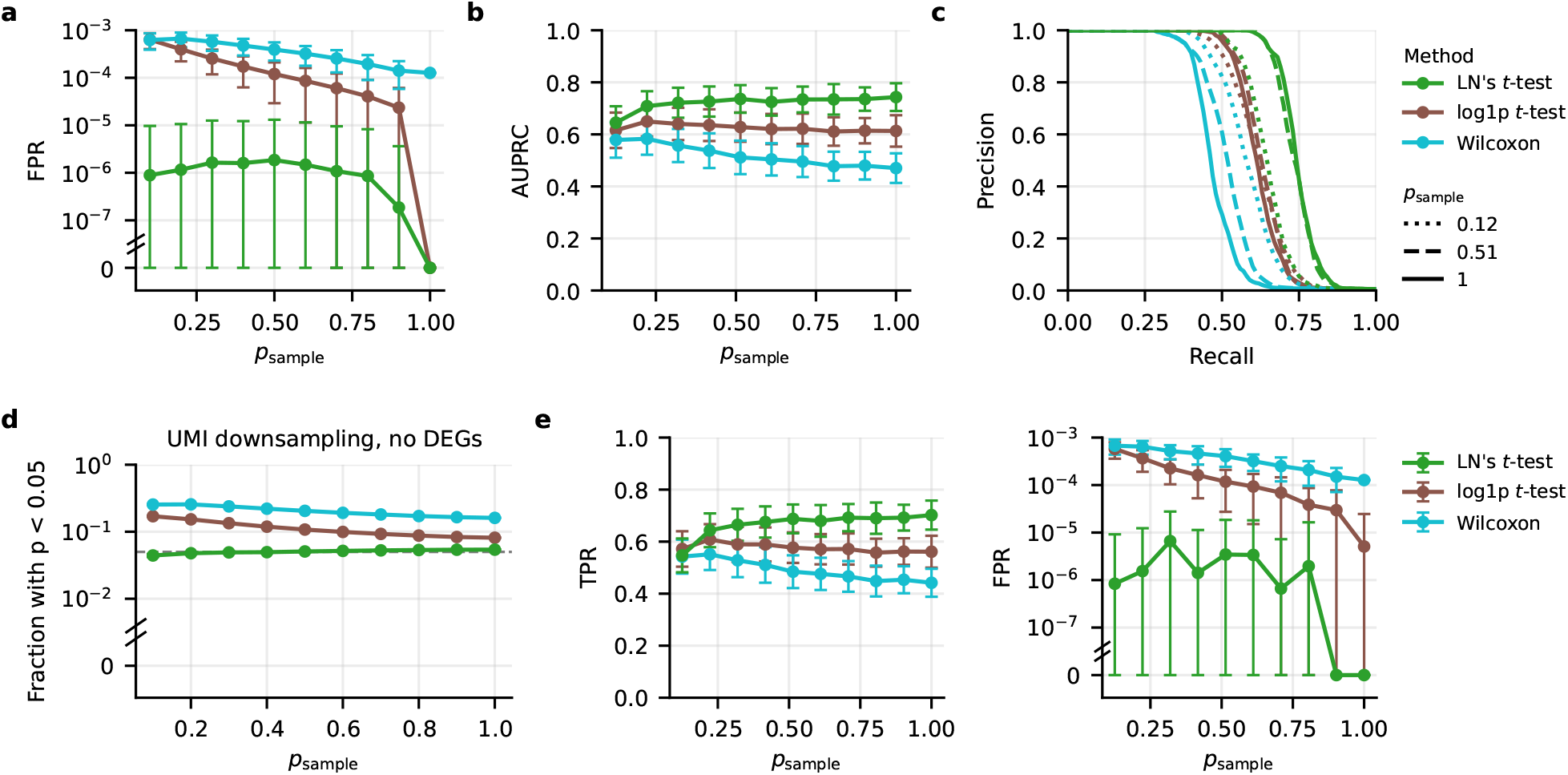
Robustness to UMI downsampling in the kidney data set. **a**, FPR without DEGs against *p*_sample_. **b**, AUPRC with simulated DEGs as in Figure 1 g. **c**, Mean precision– recall curves at *p*_sample_ = 0.12, 0.51 and 1. **d**, Fraction of genes with uncorrected *p* < 0.05 without DEGs, where a calibrated test gives 0.05 (dashed). **e**, TPR (left) and FPR (right) with simulated DEGs. Data in **a, b, d** and **e** are means and standard deviations over replicates.

We split the capsules into two groups at their median sequencing depth (Methods). LN’s *t*-test reported only 10 DEGs with all 161 capsules, compared with 1,099 for the log1p *t*-test and 3,732 for Wilcoxon, and these counts fell as the smallest capsules were removed (Supplementary Figure 4). We also performed a similar experiment focused on the effect of data sparsity, where we downsampled the UMIs of every capsule, keeping each with probability *p*_sample_, before splitting the capsules into two groups (Methods). Once again, LN remained robust to *p*_sample_ (Supplementary Figure 5 a,d), and consistently achieved the highest area under the precision–recall curve (AUPRC) in the presence of simulated DEGs (Supplementary Figure 5 b,c,e).

**Supplementary Figure 6:**
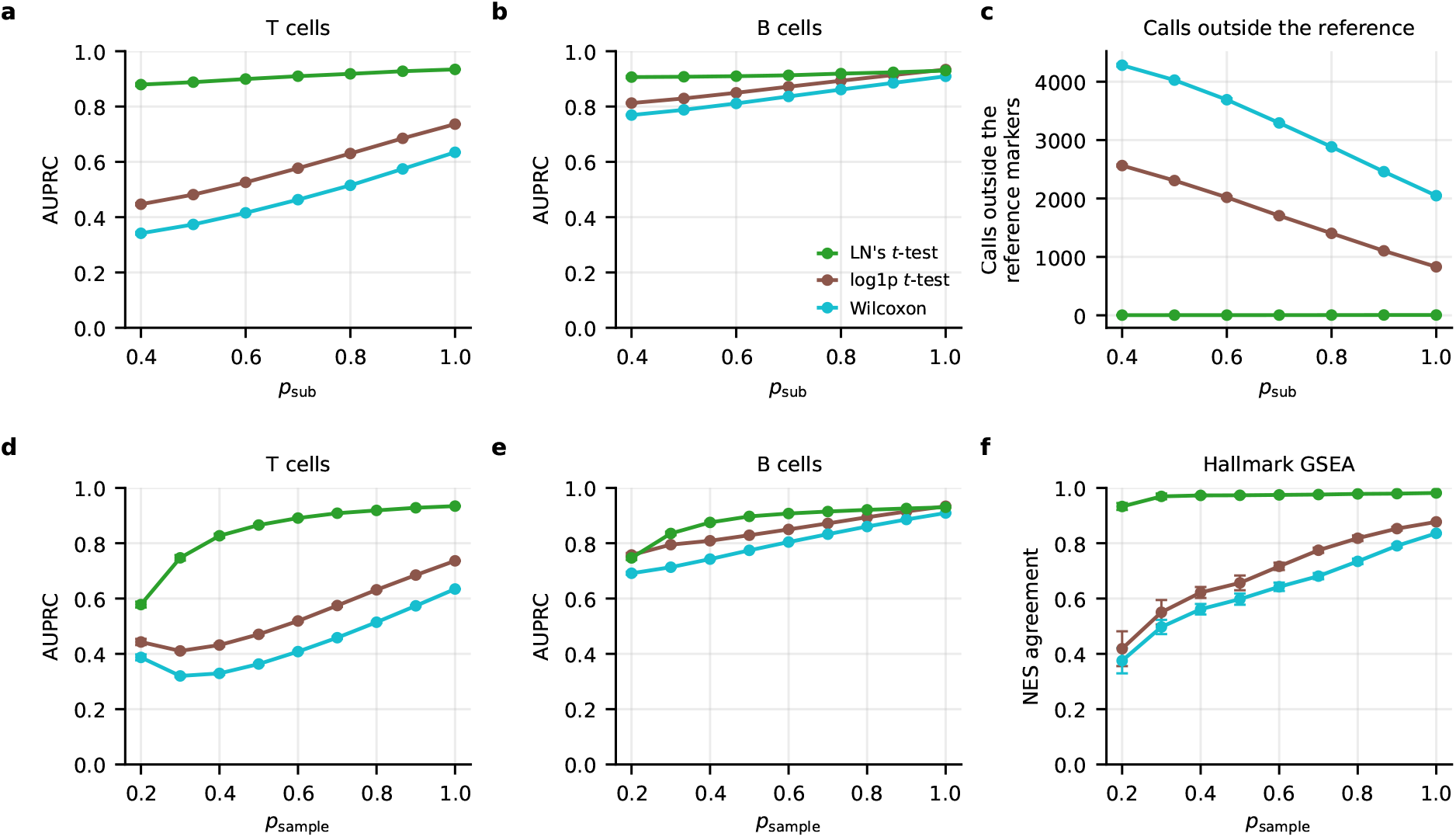
Robustness to spots and UMI downsampling in T cells vs germinal center B cells DGE on lymph node Visium HD data. **a**,**b**, AUPRC of (**a**) T-cell and (**b**) B-cell DEGs, against *p*_sub_. **c**, Number of genes each test calls outside the reference markers, against *p*_sub_. **d**,**e**, AUPRC of (**d**) T-cell and (**e**) B-cell DEGs, against *p*_sample_. **f**, Agreement of each test’s Hallmark normalized enrichment scores (NES) with the reference, against *p*_sample_. Data are means and standard deviations over replicates.

### Supplementary Results 2: Lymph node

In the lymph node experiment of Figure 2 a–c, T cells were compared with germinal center B cells in a Visium HD section, with each 2 *µ*m spot of the T cells kept with probability *p*_sub_ to lower their sequencing depth without changing their biology, and each test scored against the markers of a pseudobulk test on the full data (Methods, Supplementary Figure 6 a–c). LN recovered the T-cell markers best (Figure 2 b). For the B-cell DEGs, the baselines matched LN’s *t*-test on the full data but again performed worse as the T cells were downsampled (Supplementary Figure 6 b). We additionally downsampled UMIs instead of spots and obtained qualitatively similar results (Supplementary Figure 6 d–f). We also note that the log1p *t*-test performed better than Wilcoxon in these experiments, presumably because, like the pseudobulk reference, it compares means (of log1p-transformed counts), whereas Wilcoxon compares distributions (Supplementary Note H).

### Supplementary Results 3: PBMCs

In the clustering of Figure 2 d–f, every Leiden cluster of PBMC10k and PBMC3k, at five resolutions, was tested against all other cells (Methods), and in PBMC10k LN’s top 20 genes per cluster held the fewest housekeeping genes (Figure 2 f).

In PBMC10k, the variance imbalance of Figure 2 e (PBMC3k: Supplementary Figure 7 a) correlated with the difference in sequencing depth between the two groups involved in each test, with a weaker correlation in the smaller PBMC3k data (Supplementary Figure 7 b). In both data sets, each cluster’s median test statistic correlated with its depth ratio for the baseline methods (Pearson’s *r* of 0.63 to 0.79), but not for LN in PBMC10k (*r* = 0.00) and less so in PBMC3k (*r* = 0.44, Supplementary Figure 7 c,d). At the finer resolutions of PBMC10k, LN’s top 20 also held the largest fraction of genes that the Human Protein Atlas lists as enriched in the cluster’s cell type [10] (Supplementary Figure 7 f). In PBMC3k, which has no protein measurements, LN’s top 20 held fewer housekeeping genes and more cell-type-enriched genes than the log1p *t*-test’s at every resolution, while its advantage over Wilcoxon narrowed as the clusters became finer and reversed at the finest resolutions (Supplementary Figure 7 e,g). As the clusters became smaller, LN’s *t*-test and Wilcoxon called fewer DEGs per cluster, with LN calling the fewest at every resolution, while the log1p *t*-test consistently called more than a thousand per cluster even at the finest resolution (Extended Data Fig. 4 a,b). The growth of scanpy’s LFC with resolution (Extended Data Fig. 4 c,d) is expected, because its LFC approximation can greatly amplify small differences between transformed means (Supplementary Note I).

### Supplementary Results 4: Glioma myeloid cell atlas

In the glioma myeloid cell atlas [11] of Figure 2 g–i, each of the nine myeloid clusters (MC1–MC9, Extended Data Fig. 5 a) was tested against the other myeloid cells, and its genes, ranked by the Wilcoxon *z*-score or by LN’s score, were tested for enrichment of the Hallmark gene sets (Methods).

**Supplementary Figure 8:**
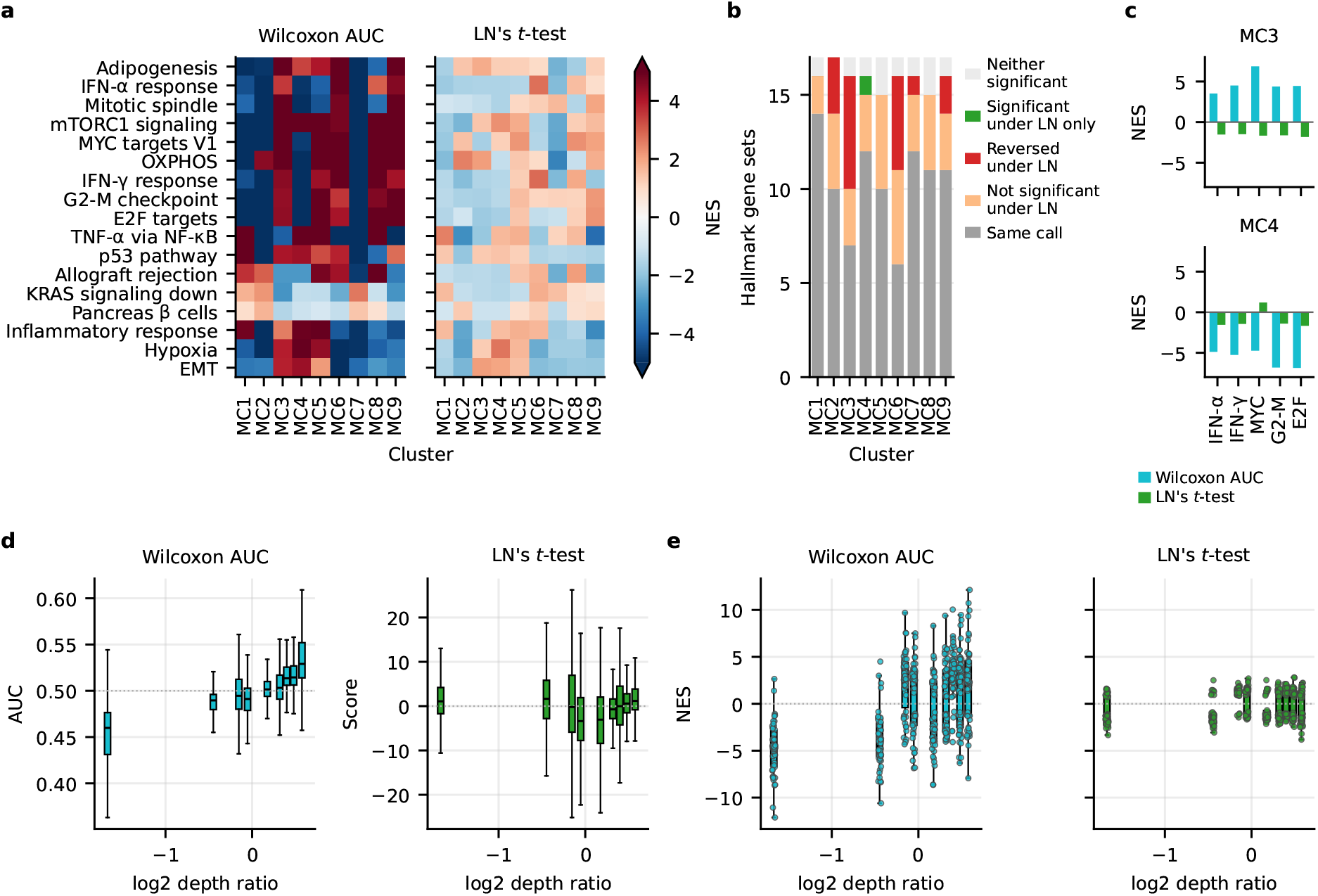
scRNA-seq data from a glioma myeloid cell atlas. **a**, NES of Hallmark gene sets for each cluster (MC1–MC9) against the other myeloid cells, with genes ranked by the Wilcoxon AUC (left) or LN’s score (right). **b**, The number of gene sets that keep their result under LN (same sign), lose it, reverse it, gain one, or are not significant under either test. **c**, NES of MC3 and MC4 in the interferon and proliferation gene sets. **d**, Wilcoxon AUC and LN’s score of expressed genes for each cluster vs depth ratios. **e**, NES of Hallmark gene sets per cluster vs log_2_ depth ratio using Wilcoxon AUC and LN’s score.

Wilcoxon called more DEGs than LN in every cluster, and the ratio grew with the depth imbalance (Extended Data Fig. 5 d–f). Wilcoxon’s differential expression scores and pathway enrichments also followed cluster depth, with the shallowest clusters depleted of most gene sets, while the deepest mostly had enrichments, whereas LN’s shifted much less with depth (Extended Data Fig. 5 g,h). The original study ranked genes by the Wilcoxon AUC, and with this procedure we reproduced 129 of the 135 enrichment directions reported in the original study (Supplementary Figure 8 a, Methods). With this ranking, MC3 was enriched and MC4 depleted in all five pathways of Figure 2 i (Supplementary Figure 8 c). We ranked by the Wilcoxon *z*-score in Figure 2 g–i because 94% of the AUCs lie between 0.45 and 0.55, which leaves the enrichment test nearly unweighted. With the AUC, the differences from LN were larger: of the Wilcoxon’s 142 significant calls, 17 were reversed and 32 not significant under LN, against 1 and 11 of 89 with the *z*-score (Supplementary Figure 8 a–c).

